# Identification of enhancer regulatory elements that direct epicardial gene expression during zebrafish heart regeneration

**DOI:** 10.1101/2021.08.23.457406

**Authors:** Yingxi Cao, Yu Xia, Joseph B. Balowski, Jianhong Ou, Lingyun Song, Alexias Safi, Timothy Curtis, Gregory E. Crawford, Kenneth D. Poss, Jingli Cao

## Abstract

The epicardium is a mesothelial tissue layer that envelops the heart. Cardiac injury activates dynamic gene expression programs in epicardial tissue, which in the case of zebrafish enables subsequent regeneration through paracrine and vascularizing effects. To identify tissue regeneration enhancer elements (TREEs) that control injury-induced epicardial gene expression during heart regeneration, we profiled transcriptomes and chromatin accessibility in epicardial cells purified from regenerating zebrafish hearts. We identified hundreds of candidate TREEs, defined by increased chromatin accessibility of non-coding elements near genes with increased expression during regeneration. Several of these candidate TREEs were incorporated into stable transgenic lines, with 5 of 6 elements directing injury-induced epicardial expression but not ontogenetic epicardial expression in hearts of larval animals. Whereas two independent TREEs linked to the gene *gnai3* showed similar functional features of gene regulation in transgenic lines, two independent *ncam1a*-linked TREEs directed distinct spatiotemporal domains of epicardial gene expression. Thus, multiple TREEs linked to a regeneration gene can possess either matching or complementary regulatory controls. Our study provides a new resource and principles for understanding the regulation of epicardial genetic programs during heart regeneration.

## INTRODUCTION

The zebrafish heart is capable of complete or near complete regeneration after injury, based on proliferation of spared cardiomyocytes (CMs) (Poss et al., 2002). The pro-regenerative environment provided by non-muscle cells such as the epicardium, endocardium, vasculature, and immune cells contributes to this potential (Cao and Poss, 2018; Gemberling et al., 2015; Gonzalez-Rosa et al., 2012; Hui et al., 2017; Karra et al., 2018; Kikuchi et al., 2011; Lepilina et al., 2006; Masters and Riley, 2014; Wang et al., 2013). For example, genetic ablation of the epicardium, a thin mesothelial layer that envelops all vertebrate hearts, blocks heart muscle regeneration and coronary angiogenesis in zebrafish (Wang et al., 2015). Key, developmentally potent genes like retinaldehyde dehydrogenase 2 (*raldh2*), T-box transcription factor 18 (*tbx18*), fibronectin 1 (*fn1*), and neuregulin 1 (*nrg1*) are induced in epicardial tissue upon cardiac injury, first organ-wide and then resolving to the site of trauma (Figure 1A), in a phenomenon known as “epicardial activation” (Gemberling et al., 2015; Lepilina et al., 2006; Wang et al., 2013). Understanding the gene expression responses to injury that define epicardial activation can illuminate defining aspects of heart regeneration (Cao and Poss, 2018).

**Figure 1.**
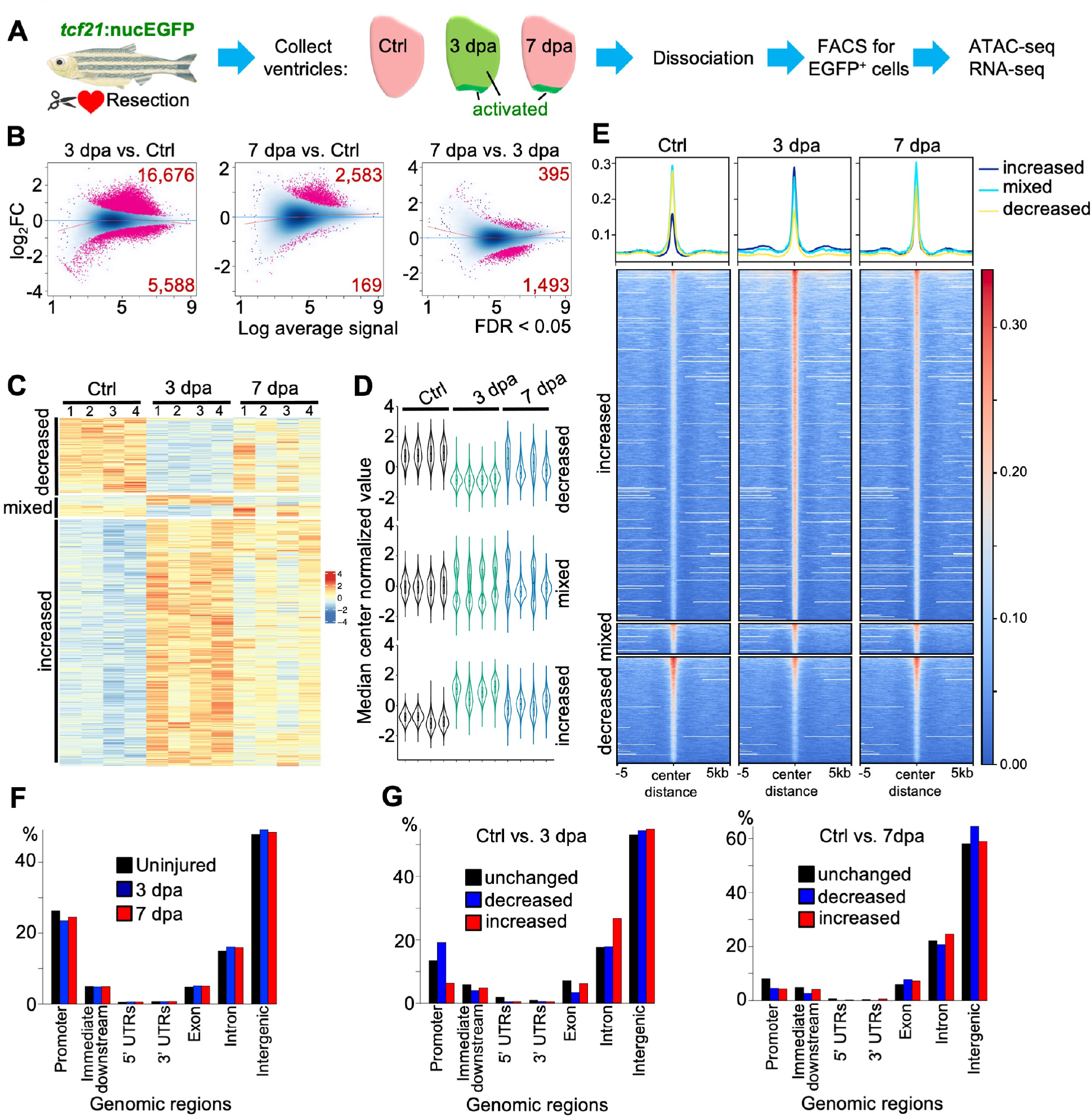
ATAC-seq analysis reveals dynamic chromatin accessibility in the regenerating epicardium. **(A)** Schematic for experiment design. Partial resection injuries were performed in *tcf21:nucEGFP* animals. Ventricles were collected at 3 or 7 dpa, as were those of uninjured clutchmates (Ctrl). Epicardial activation regions are labeled in green in the cartoons. Ventricles were dissociated, EGFP^+^ epicardial cells were isolated by FACS, and bulk RNA-seq and ATAC-seq were performed. **(B)** MA plots of Log_2_ fold changes over average normalized ATAC-seq signals. Red dots denote peaks with significantly changed chromatin accessibility with the numbers of differential peaks labeled in the corners (FDR < 0.05). **(C)** Heat map of differential chromatin accessibility across 3 groups with 4 replicates each. Three clusters (increased, mixed, and decreased) were shown on the left. Increased, peaks with increased accessibility in both 3 and 7 dpa samples. Decreased, peaks with decreased chromatin accessibility in both 3 and 7 dpa samples comparing with the uninjured control. Mixed, peaks with different trends in 3 and 7 dpa samples, comparing with the control. **(D)** Violin plots of differential peaks of three clusters across samples, showing the distribution pattern of chromatin accessibility across samples. **(E)** Heat map shows the signals of ATAC-seq within ± 5 kb from peak centers. The blue, cyan and yellow lines at the top represent mean read densities of the corresponding ATAC-seq peaks at increased, mixed, and decreased chromatin accessibility, respectively **(F)** Genomic distributions of all ATAC-seq peaks in three groups. **(G)** Genomic distributions of the differential and unchanged regions in pair-wised comparisons (left, 3 dpa vs. Ctrl; right, 7 dpa vs. Ctrl).

Enhancers are a class of cis-regulatory elements that help orchestrate gene expression during animal development and in response to environmental changes (Pennacchio et al., 2013). Kang et al. first reported a short non-coding DNA sequence upstream of the gene *leptin b* (*lepb*) that can direct expression in zebrafish hearts and fins upon injury and during regeneration, referring to these context-preferential sequences as tissue regeneration enhancer elements (TREEs). TREEs can be identified by comparing profiles of chromatin structure or decorations from uninjured and regenerating tissues. From these profiles, sequences near genes that increase RNA levels during regeneration, and in which increased marks of activated enhancers are evident in the regeneration contexts, represent candidate TREEs. Generation of transgenic animals and/or mutant animals is essential to validate TREEs, which have been described and validated in many contexts including zebrafish hearts, zebrafish and killifish fins, and *Drosophila* imaginal discs (Begeman et al., 2020; Goldman et al., 2017; Harris et al., 2016; Kang et al., 2016; Pfefferli and Jazwinska, 2017; Wang et al., 2020). The discovery of specific regulatory sequences that underlie regeneration programs can reveal candidate upstream and downstream factors in regeneration, while also providing tools to manipulate regeneration (Chen et al., 2020; Kang et al., 2016; Sugimoto et al., 2017; van Duijvenboden et al., 2019).

Initial studies have made it clear that different cell populations engage distinct compendia of TREEs during their respective regenerative responses (Goldman et al., 2017; Kang et al., 2016; Lee et al., 2020; Thompson et al., 2020). Here, to elucidate candidate TREEs responsible for epicardial gene expression responses, we profiled the chromatin accessibility of epicardial cells during heart regeneration in zebrafish by ATAC-seq (Buenrostro et al., 2013), validating several of these TREEs using stable transgenic reporter lines. Our study provides a resource of gene regulatory changes in epicardial cells during heart regeneration and reveals new concepts in TREE-based gene regulation.

## RESULTS AND DISCUSSION

### ATAC-seq analysis of epicardial chromatin structure during heart regeneration

To identify candidate TREEs that direct epicardial gene expression, we profiled transcriptomes and whole-genome chromatin accessibility by bulk RNA-seq and ATAC-seq from purified epicardial cells. We postulated that these datasets would reveal areas of active regulation, including enhancer elements linked to genes involved in epicardial responses to injury and regeneration. To elicit a strong epicardial injury response, we performed ventricular resection injuries on *tcf21:nucEGFP* animals and collected ventricles 3 and 7 days later (dpa, days post-amputation) together with uninjured ventricles (Ctrl) for isolation of *tcf21*^+^ cells (Figure 1A). These two time-points represent a stage of organ-wide epicardial activation (3 dpa) and injury site-restricted activation (7 dpa, Figure 1A) (Lepilina et al., 2006). From four biological replicates, we identified 315k open chromatin sites on average in each experimental group (Table S1). In 3 dpa samples, 16,676 sites displayed increased chromatin accessibility compared to samples from uninjured hearts, whereas 5,588 had decreased accessibility (Figure 1B and Table S2, false discovery rate (FDR) < 0.05). By 7 dpa, chromatin accessibility appeared to largely normalize: 2,583 sites had significantly increased chromatin accessibility, and 169 sites had decreased accessibility (Figure 1B). Clustering of these differential sites across samples derived three clusters with distinct changes in chromatin accessibility. The first cluster displayed increased accessibility during regeneration in both 3 and 7 dpa samples (Figure 1C-E, increased), while the second cluster showed reduced accessibility (Figure 1C-E, decreased). The last cluster has mixed trends (increased or decreased) in 3 and 7 dpa samples, with the 7 dpa sample more similar to the control (Figure 1C-E, mixed). We next analyzed peak distribution by genomic regions. In all 3 experimental groups, about 25% of the peaks resided in promoters, ∼15% in introns, ∼5% in exons, and ∼50% are intergenic. The rest are within the untranslated regions (UTRs) and immediate downstream regions (Figure 1F). The increased regions at 3 dpa (vs. Ctrl) mostly resided in the intergenic (∼55%) and intronic regions (∼25%) (Figure 1G). Together, these results suggest that there is substantial chromatin remodeling in epicardial cells after injury, and this remodeling is more extreme at 3 dpa than 7 dpa.

To more closely examine these data for active regulatory elements, we integrated our ATAC-seq dataset with the published histone H3K27Ac (H3 acetylation at lysine 27) signature captured from regenerating zebrafish ventricles (Kang et al., 2016). This dataset comprises profiles of two biological replicates of ventricles regenerating after partial genetic ablation of CMs (regenerating), and uninjured ventricles (control). We examined trends of differential ATAC peaks, finding that regions with either increased or decreased accessibility during regeneration bear H3K27Ac signatures in whole-ventricle samples (Figure 2A). However, only those regions with increased accessibility correlate well with a signature of increased H3K27Ac marks in regenerating samples (Figure 2B, regenerating/control ratio > 1). By contrast, those regions decreasing do not show changes in the H3K27Ac signature (Figure 2B, bottom, ratio = 1). As an example, Figure 2C shows the genomic region comprising Wilms tumor 1 transcription factor b (*wt1b*), a key epicardial transcription factor induced by injury in zebrafish (Kikuchi et al., 2011). Four peaks upstream (-13 kb, -19 kb, -20 kb, and -24 kb) of the transcription start site (TSS) demonstrated increased accessibility at both 3 and 7 dpa. Three of these peaks have increased H3K27Ac marks in the regenerating ventricle dataset (peaks 1-3; additional examples in Figure S1). This analysis implicates regions of DNA with increased chromatin accessibility in epicardial cells during heart regeneration as candidate TREEs.

**Figure 2.**
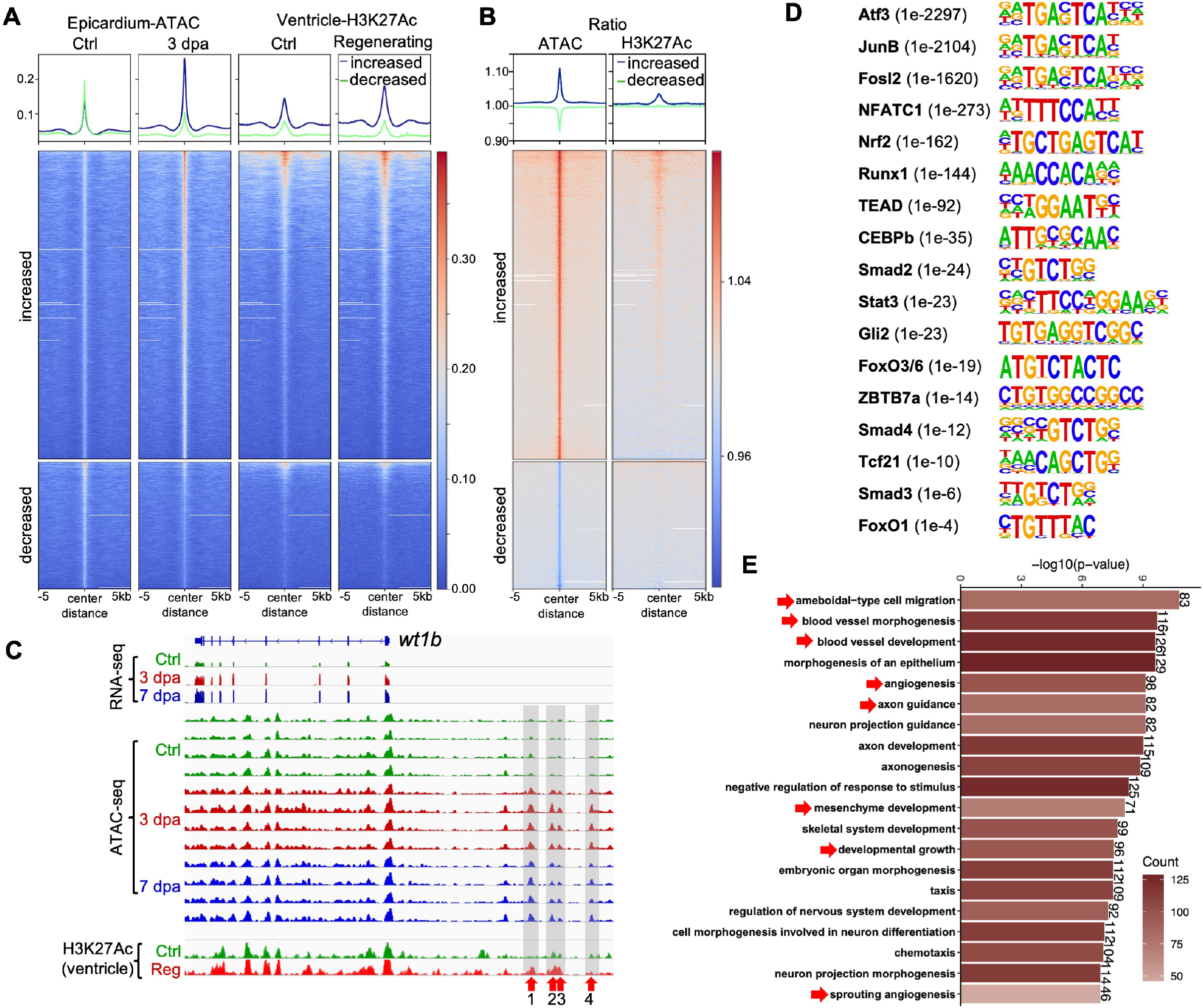
Chromatin accessibility indicates regulatory programs of regenerating epicardium. **(A)** Heat map with signals of ATAC-seq and Chip-seq within ± 5 kb from peak centers. The blue and green lines at the top represent mean read densities of the corresponding ATAC-seq peaks with increased and decreased chromatin accessibility (3 dpa vs. Ctrl), respectively. **(B)** Heat map indicates ratios of the signals shown in (A). 3 dpa/Ctrl for ATAC-seq (left), and Regenerating/Ctrl for Chip-seq (right). **(C)** Browser tracks of the genomic region comprising gene *wt1b* showing the transcripts and chromatin accessibility profiles in the epicardium across replicates from Ctrl, 3 dpa, and 7 dpa samples. The whole-ventricle H3K27Ac profile of the uninjured (Ctrl) and regenerating (Reg) hearts is shown at the bottom. Gray boxes, red arrows, and numbers indicate ATAC-seq peaks with increased accessibility during regeneration. **(D)** Enriched motifs in the open chromatin regions that gained accessibility at 3 dpa. P-value for each enriched TF is shown in the bracket. **(E)** Barplot showing the 20 most enriched GO biological function terms obtained from peaks with increased chromatin accessibility in 3 dpa samples (FDR < 0.01). The full list is shown in Table S3.

### Motif and pathway enrichment analysis of regions with context-specific accessibility changes

Recurrent consensus motifs that gain active enhancer marks are likely to contain binding sites for transcription factors (TFs). To identify candidate transcriptional regulators active in epicardial cells during heart regeneration, we assayed for enriched nucleotide motifs within regions gaining accessibility at 3 dpa (vs. Ctrl) using HOMER (Heinz et al., 2010). Among the top hits are binding motifs of the TF activator protein-1 (AP-1) subunits, such as Atf3 (activating transcription factor 3), JunB, and Fos (Figure 2D), present in about half of the analyzed regions. The AP-1 complex was recently implicated in the control of CM gene expression during zebrafish heart regeneration (Beisaw et al., 2020). However, AP-1 motifs are common within regulatory sequences (Umer et al., 2019), and thus a requirement for these motifs might not reflect specificity for regeneration-related gene expression. Other enriched motifs highlight pathways and TFs known to regulate epicardium development and/or regeneration: Tcf21 (Hu et al., 2020), Hippo/Yap pathway (TEA domain family members, TEADs) (Xiao et al., 2018), C/EBPb (CCAAT/enhancer-binding protein b)(Huang et al., 2012), Tgfβ pathway (Smad2/3/4) (Chablais and Jazwinska, 2012), Hedgehog pathway (GLI family zinc finger 2, Gli2) (Choi et al., 2013; Sugimoto et al., 2017; Wang et al., 2015). Additional implicated TFs that have not been connected to epicardial functions include NFATC1 (Nuclear Factor of Activated T Cells 1), Nrf2 (Nuclear factor-E2-related factor 2), Stat3, FoxO1 (Forkhead box O1), FoxO3, FoxO6, and ZBTB7A (Zinc Finger and BTB Domain Containing 7A) (Figure 2D).

For an overview of biological functions of genes linked to dynamic chromatin regions at 3 dpa, we performed Gene Ontology (GO) enrichment analysis. Comparing to epicardial cell samples from uninjured hearts, we found several enriched pathways, including Tgfβ signaling pathway and FoxO signaling pathway, that match the motif analysis results (Table S3). The FoxO pathway regulates many cellular physiological processes such as apoptosis, cell cycle, metabolism, and oxidative stress resistance, by acting downstream of growth factors, insulin, glucose, Tgfβ, and other stimulators (Lu and Huang, 2011; Nakae et al., 2001; Wang et al., 2014). Other enriched pathways include cellular senescence and adherens junctions. It was recently reported that p53 induces senescence in mouse epicardial cells upon heart injury, and induced cellular senescence promotes neonatal heart regeneration (Feng et al., 2019; Sarig et al., 2019). The abovementioned transcription factor Nrf2 and Stat3 are key regulators of cellular senescence (Yan et al., 2021). The enriched processes of blood vessel development and angiogenesis are consistent with the role of the epicardium in supporting revascularization during regeneration (Marin-Juez et al., 2019; Wang et al., 2015). Biological processes such as mesenchymal cell development and differentiation, stem cell development and differentiation, implicate the progenitor feature of the epicardial cells. Other enriched processes include heart morphogenesis, apoptotic signaling pathway, cell migration involved in heart development, and, unexpectedly, axon development-related processes (Figure 2E and Table S3). Enriched molecular functions further emphasized ligand-receptor activities, extracellular matrix binding, and SMAD binding (Table S3).

### Direct tests of candidate regulatory elements for enhancer activity during regeneration

To prioritize a list of candidates for functional validation of enhancer activity, we first examined the top 20 distal regions with the highest fold increases in accessibility during heart regeneration (Figure 3A). These peaks were assigned to nearest genes, which included fibronectin 1a (*fn1a*) - an epicardially expressed gene that is required for zebrafish heart regeneration (Wang et al., 2013), neural cell adhesion molecule 1a (*ncam1a*) - a gene involved in axon development (Siles et al., 2018) (listed by the enrichment annotation results in Figure 2E and Table S3), repulsive guidance molecule BMP co-receptor b (*rgmb*) – encoding a TGF-β superfamily signaling component that participates in neuronal development (Liu et al., 2016; Samad et al., 2005) (Table S3), and guanine nucleotide binding protein, alpha inhibiting activity polypeptide 3 (*gnai3*) – encoding a G protein. We next combined the bulk RNA-seq and ATAC-seq datasets and identified 3,026 ATAC-peaks with their related 1,008 genes and RNA levels, with both features increased at 3 dpa (Figure 3B, red dots; Figure 3C; Tables S4, S5). *fn1a*, *rgmb*, and *gnai3* are among the list (Figure 3B). Lastly, we looked for conserved regions by comparing our dataset with the published zebrafish Conserved Non-genic Elements (zCNEs) database, which comprises conserved regions from fish to human (Hiller et al., 2013). We identified 886 ATAC-seq peaks that contain conserved regions and gain accessibility at 3 dpa (Figure 3D and Table S6), including the top differential peaks assigned to *ncam1a* (Figure 3E, Enhancer 2 or E2). As shown in Figure 3E, four emerging ATAC-seq regions reside in the intronic regions of *ncam1a*, which increase accessibility at both 3 and 7 dpa. *ncam1a-E2* (+181 kb) contains conserved sequences and displayed strong enrichment with histone H3K27Ac mark in samples of whole regenerating ventricles. *ncam1a-E4* (+110 kb) also has a significantly enriched histone H3H27Ac signature. These regions received highest priority for functional tests in transgenic lines.

**Figure 3.**
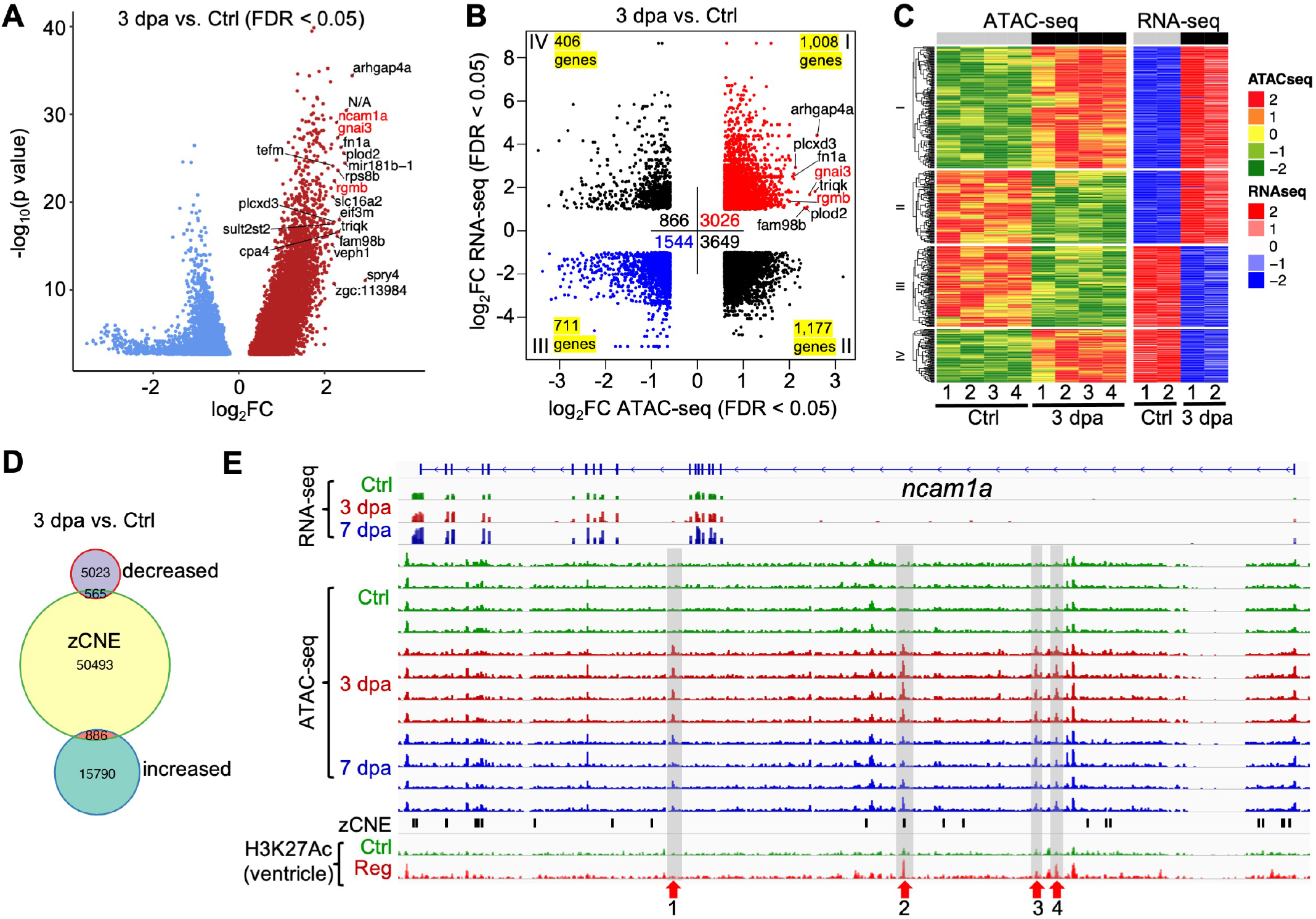
Candidate regulatory elements for enhancer activity during regeneration. **(A)** Volcano plot of differential ATAC-seq peaks in 3 dpa versus uninjured (Ctrl) samples. The top 20 upregulated peaks with the most fold changes are marked with the annotated genes. *ncam1a*, *gnai3* and *rgmb* are highlighted in red. N/A, no gene is annotated to the peak. **(B)** Dot plot of differential ATAC-seq peaks linked to nearby differential transcripts in 3 dpa versus uninjured (Ctrl) samples. Each dot indicates an individual ATAC-seq peak and is counted into the total peak numbers of each quadrant. The number of unique genes in each quadrant is labeled in the corner. The most differential peaks of a few genes are marked with *gnai3* and *rgmb* highlighted in red. **(C)** Heat map of differential transcripts in 3 dpa versus uninjured (Ctrl) linked to nearby differentially accessible chromatin regions. **(D)** Venn diagram comparison of differentially regulated ATAC-seq peaks in 3 dpa versus uninjured compared with the zCNE list. **(E)** Browser tracks of the genomic region near gene *ncam1a* showing the transcripts and chromatin accessibility profiles in the epicardium. The whole-ventricle H3K27Ac profile of the uninjured (Ctrl) and regenerating (Reg) heart is shown at the bottom. zCNE sites are depicted as short black bars. Gray boxes, red arrows, and numbers indicate candidate TREEs.

#### A rgmb-linked TREE directs injury-induced epicardial gene expression

Rgmb, a TGF-β superfamily signaling component, has been reported to bind to BMP2 or BMP4 as a co-receptor, to pattern the developing nervous system or to inhibit renal cyst development (Liu et al., 2016; Samad et al., 2005). Although no function of *rgmb* in heart regeneration has been reported, *bmp2b* overexpression enhances CM proliferation after cardiac injury in zebrafish (Wu et al., 2016). *rgmb* RNA levels increase at 3 dpa (log_2_FC = 1.74, Padj (adjusted P-value) = 7.42E-12) and 7 dpa (log_2_FC = 1.38, Padj= 1.92E-06) in our datasets (Table S4). We searched for *rgmb* in the Zebrafish Regeneration Database that comprises published transcriptome datasets of heart, fin, or spinal cord regeneration (http://zfregeneration.org) (Nieto-Arellano and Sanchez-Iranzo, 2019), finding that *rgmb* RNA levels increase during heart or spinal cord regeneration, consistent with a pro-regenerative function in different tissues (Figure S2). Two putative enhancers (*rgmb-E1*, -68 kb; *rgmb-E2*, -71 kb) upstream of the TSS were identified from our analyses, with *rgmb-E1* showing enriched H3K27Ac marks in whole-ventricle samples (Figure 4A). To test the efficacy of these candidate enhancers, we subcloned each regulatory region upstream of a *c-fos* minimal promoter and *EGFP* cassette (Figure 4B). Without a regulatory sequence, this cassette has minimal expression in embryos and adult hearts, even after cardiac injury (Goldman et al., 2017). Multiple stable lines were established. We found that *rgmb-E1* sequences direct strong EGFP expression in F1 embryos, including heart, eye, and notochord expression (Figure 4B), which resemble the published *in situ* hybridization results (Figure 4C, adapted from ZFIN) (Thisse et al., 2008). Cardiac expression in the 6 dpf (days post-fertilization) larvae is restricted to the myocardium but not the epicardium (Figure 4D). In adult hearts, injury-induced *rgmb* transcripts were detected on the ventricular surface (Figure 4E). For the *rgmb-E1* reporter lines, EGFP expression in adult epicardial cells (*tcf21*:H2A-mCherry+) was only detectable after a heart injury, both locally and distal to the injury site (Figure 4F-G), suggesting that *rgmb-E1* is an epicardial TREE. By contrast, 3 stable lines of *rgmb-E2:EGFP* were identified without consistent EGFP expression in larval or adult hearts (Figure 4H-I). These results indicate that, whereas *rgmb-E1* is sufficient to direct injury-induced gene expression, *rgmb-E2* on its own is inadequate.

**Figure 4.**
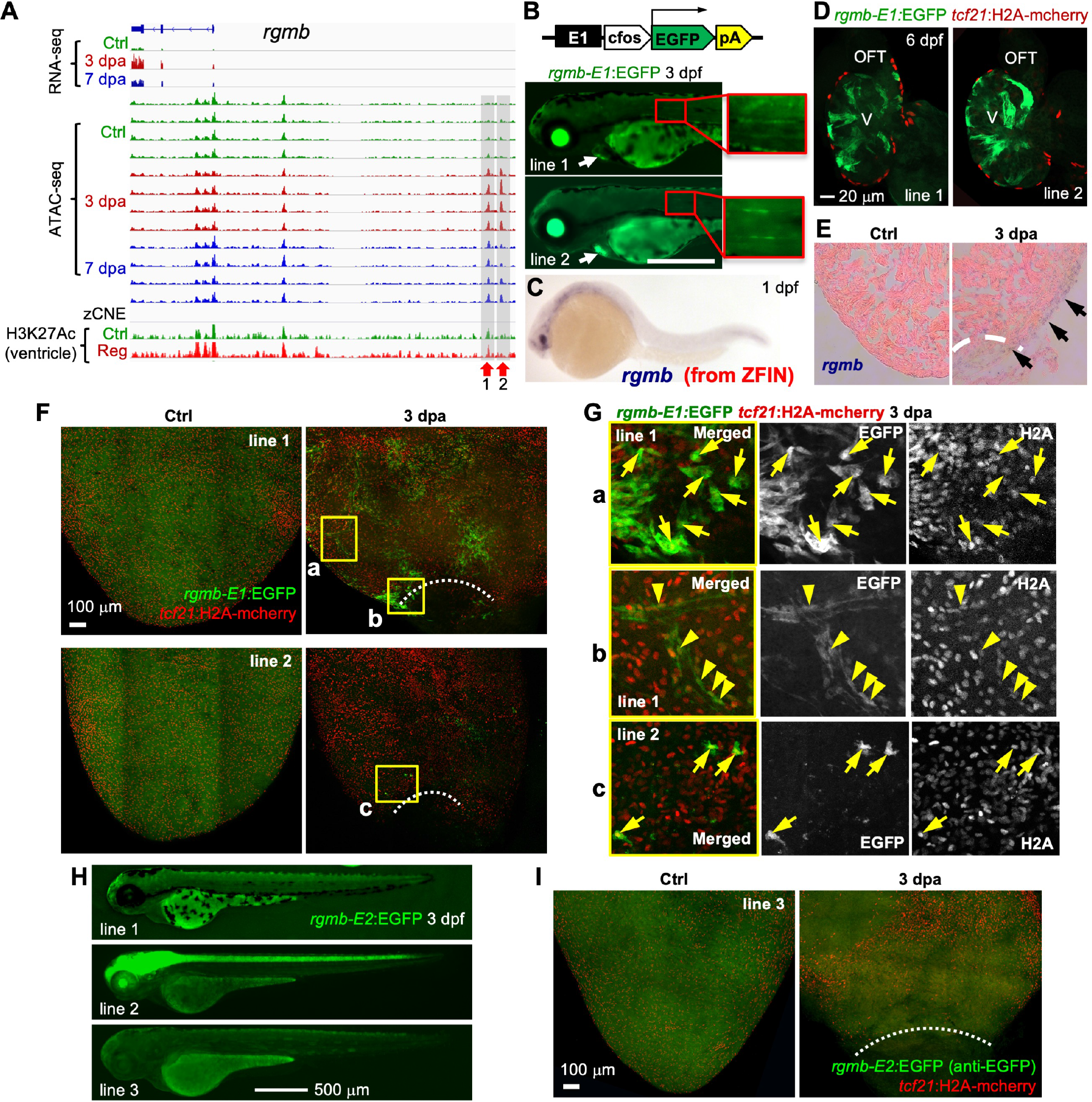
A *rgmb*-linked TREE directs injury-induced epicardial gene expression. **(A)** Browser tracks of the genomic region near gene *rgmb* showing the transcripts and chromatin accessibility profiles in the epicardium. The whole-ventricle H3K27Ac profile of the uninjured (Ctrl) and regenerating (Reg) heart is shown at the bottom. Gray boxes, red arrows, and numbers indicate candidate TREEs. **(B)** Top: the *rgmb-E1:EGFP* reporter construct. Bottom: larval expression of *rgmb-E1:EGFP* lines at 3 dpf. Scale bar, 500 μm. **(C)** Whole-mount *in situ* hybridization showing *rgmb* expression in a 1 dpf embryo. Adapted from ZFIN (Thisse et al., 2008). **(D)** Optical section images of 6 dpf hearts (whole-mounted) showing EGFP expression in the muscle. *tcf21*:H2A-mCherry (red) labels the epicardial cells. V, ventricle. Scale bar, 20 μm. **(E)** *In situ* hybridization results showing *rgmb* expression on the ventricular surface (arrows) at 3 dpa. **(F)** Whole-mount images of the ventricular surface showing expressions of the *rgmb-E1:EGFP* reporter lines (green) in uninjured (Ctrl) and 3 dpa samples. *tcf21*:H2A-mCherry (red) labels the epicardial cells. White dashed lines indicate the injury sites. The framed regions are enlarged to show details in (G). Scale bar, 100 μm. **(G)** Single-channel images are shown in grayscale. Arrows and arrowheads indicate representative EGFP^+^mCherry^+^ cells. Arrowheads in (b) point to double-positive perivascular cells. **(H)** Larval expression of *rgmb-E2:EGFP* lines. Scale bar, 500 μm. **(I)** Whole-mount images of the ventricular surface showing no EGFP induction (with an anti-EGFP antibody staining) in the 3 dpa sample carrying the *rgmb-E2:EGFP* reporter. *tcf21*:H2A-mCherry (red) labels the epicardial cells. White dashed lines indicate the injury sites. Scale bar, 100 μm.

#### Distinct gnai3-linked TREEs direct similar injury-induced epicardial gene expression

Gnai3 is a G protein that binds to G protein-coupled receptors (GPCRs) to regulate various transmembrane signaling pathways (Syrovatkina et al., 2016). The Zebrafish Regeneration Database reports increased RNA levels of *gnai3* during heart, fin, and spinal cord regeneration, suggesting functions in multiple regeneration contexts (Figure S3) (Nieto-Arellano and Sanchez-Iranzo, 2019). In our datasets, *gnai3* RNA is increased at 3 dpa (log_2_FC = 2.08, Padj = 6.43E-11) and 7 dpa (log_2_FC = 0.85, Padj = 0.023) (Table S4). We identified two putative enhancers within intron 1 (*gnai3-E1*, +8.8 kb) and intron 4 (*gnai3-E2*, +21 kb), both of which have H3K27Ac marks in the whole-ventricle profile (Figure 5A). Three stable lines for *gnai3-E1* and 4 lines for *gnai3-E2* were established. Each *gnai3-E1:EGFP* line displayed whole-body larval EGFP expression without clear specificity (Figure 5B). No EGFP expression was noticed in larval epicardium by 6 dpf (Figure 5C). In adult hearts from each of 3 lines, we consistently observed a small population of EGFP^+^; *tcf21:*H2A-mCherry^+^ cells on the ventricular surface after injury (Figure 5D, 3 dpa, based on anti-EGFP antibody staining).

**Figure 5.**
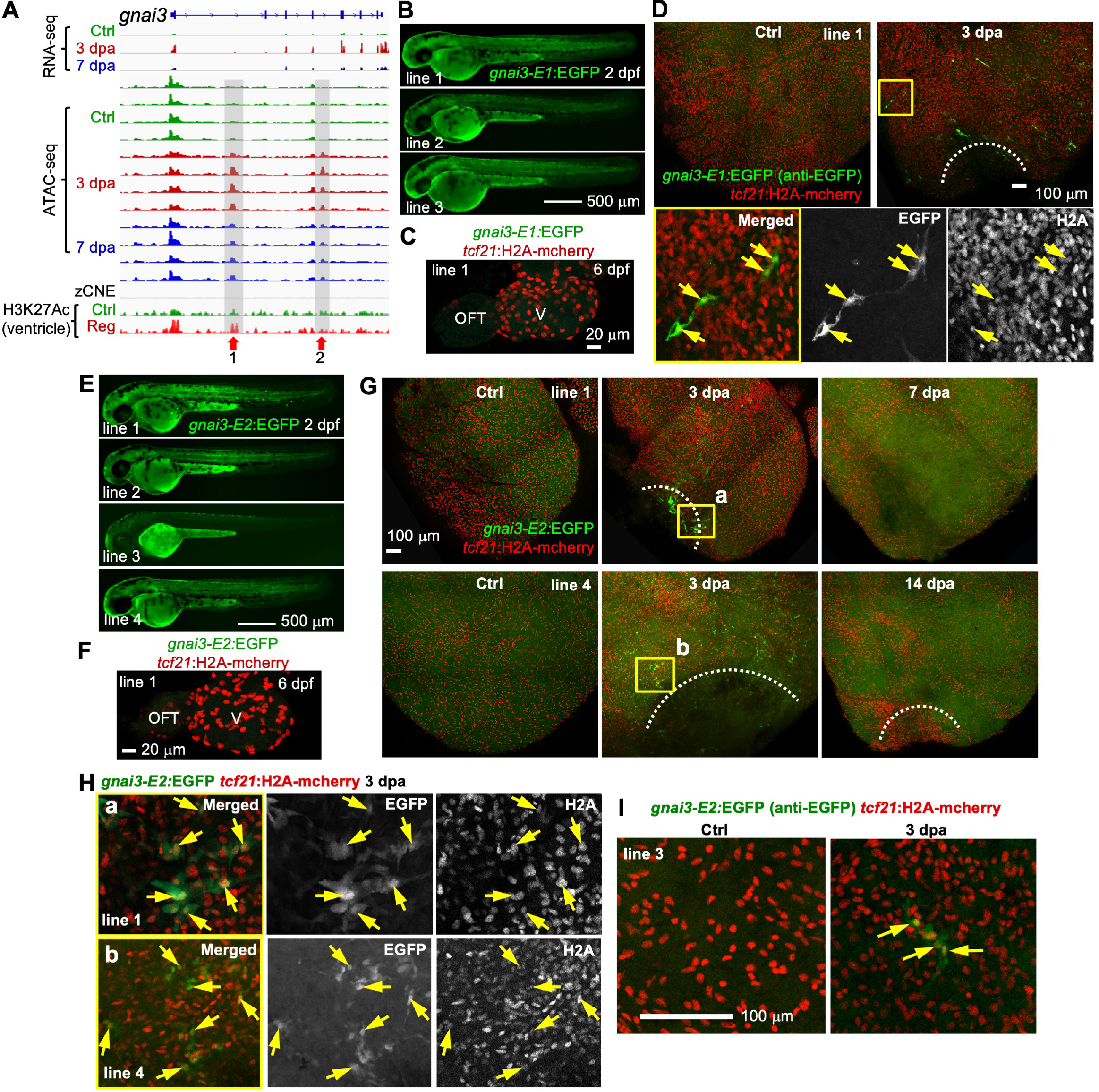
Distinct *gnai3*-linked TREEs direct similar injury-induced epicardial gene expression. **(A)** Browser tracks of the genomic region near gene *ncam1a* showing the transcripts and chromatin accessibility profiles in the epicardium, the zCNE sites, and the whole-ventricle H3K27Ac profile. Gray boxes, red arrows, and numbers indicate candidate TREEs. **(B)** Larval expression of three *gnai2-E1:EGFP*reporter lines. Scale bar, 500 μm. **(C)** Whole-mount image of a 6 dpf heart showing no GFP expression. *tcf21*:H2A-mCherry (red) labels the epicardial cells. V, ventricle. Scale bar, 20 μm. **(D)** Whole-mount images of the ventricular surface showing *gnai3-E1:EGFP* expression in *tcf21*:H2A-mCherry^+^ cells. Close-up view of the framed region is displayed at the bottom with single-channel images shown in grayscale. White dashed line indicates the injury sites. Arrows denote representative EGFP^+^mCherry^+^ cells. Scale bar, 100 μm. **(E)** Larval expression of four *gnai2-E2:EGFP* reporter lines. Scale bar, 500 μm. **(F)** Whole-mount image of a 6 dpf heart showing no EGFP expression. *tcf21*:H2A-mCherry (red) labels the epicardial cells. V, ventricle. Scale bar, 20 μm. **(G)** Whole-mount images of the ventricular surface showing expressions of the *gnai3-E2:EGFP* lines 1 and 4 (green) in uninjured (Ctrl) and 3, 7, and 14 dpa samples. *tcf21*:H2A-mCherry (red) labels the epicardial cells. White dashed lines indicate the injury sites. The framed regions are enlarged to show details in (H). Scale bar, 100 μm. **(H)** Single-channel images are shown in grayscale. Arrows denote representative EGFP^+^mCherry^+^ cells. **(I)** Whole-mount images showing expressions of the *gnai3-E2:EGFP* reporter line 3. A few EGFP^+^; *tcf21*:H2A-mCherry^+^ cells were seen upon injury (arrows). Scale bar, 100 μm.

*gnai3-E2:EGFP* is expressed in the notochord and regions of skin consistently in several lines, in addition to the weak whole-body expression (Figure 5E). Similar to *gnai3-E1:EGFP,* no embryonic epicardial expression was seen for *gnai3-E2:EGFP* lines by 6 dpf (Figure 5F). In adults, images of lines 1 and 4 of *gnai3-E2:EGFP* demonstrated injury-induced expression in epicardial cells around the injury site at 3 dpa. These expression patterns are diminished by 7 or 14 dpa (Figure 5G-H), mimicking the reduced RNA levels of *gnai3* at 7 dpa (Figure 5A). The injury induced EGFP expression of line 3 is relatively weak but restricted to *tcf21*^+^ cells (based on anti-EGFP antibody staining, Figure 5I). These results suggest that *gnai3-E1* and *E2* are epicardial TREEs, and their similar expression dynamics indicate how the expression pattern of one gene during regeneration may receive similar regulatory instructions from two distinct TREEs. Although genes are commonly regulated by multiple enhancers with redundant activities in developmental contexts (reviewed in (Kvon et al., 2021)), our finding now extends this concept to tissue regeneration.

#### Distinct ncam1a-linked TREEs direct different domains of epicardial gene expression during heart regeneration

The human homolog *NCAM1* encodes a cell adhesion protein of the immunoglobulin superfamily that regulates cell-cell and cell-matrix interactions during development and differentiation processes (Duncan et al., 2021; Siles et al., 2018). The Zebrafish Regeneration Database recorded that *ncam1a* RNA levels increase during heart, fin, or spinal cord regeneration (Figure S4). We identified two stable lines for *ncam1a-E2:EGFP*. In 3 dpf larvae, line 1 directs strong EGFP expression in the eye, brain, spinal cord, and notochord, resembling *ncam1a* expression in the embryo (Figure 6A (adapted from ZFIN) (Thisse et al., 2008), 6B). Line 2 has dimmer expression, but EGFP signals are still visible in these tissues (Figure 6B). In addition, whole-mount images of dissected 6 dpf hearts indicating EGFP expressions primarily in the outflow tract and atrioventricular valves, but not in *tcf21*^+^ epicardial cells (Figure 6C). After resection of the adult ventricle, we observed *ncam1a* expression in the ventricular surface around the injury site at 3 dpa, suggesting an epicardial expression pattern (Figure 6D). Similarly, images of both *ncam1a-E2:EGFP* lines demonstrated injury-induced EGFP expression around the wound at 3 and 7 dpa in *tcf21*:H2A-mCherry^+^ cells (Figure 6E-I), including the perivascular cells (Figure 6F, arrowheads). By 14 dpa, EGFP expression is much weaker but broadly distributed on the entire ventricular surface and primarily in the perivascular cells (Figure 6E, 6Fd-e). EGFP was almost undetectable at 30 dpa (Figure 6E). Although line 2 has lower embryonic expression than line 1, its adult heart expression pattern is consistent with line 1 (Figure 6G-I and data not shown).

**Figure 6.**
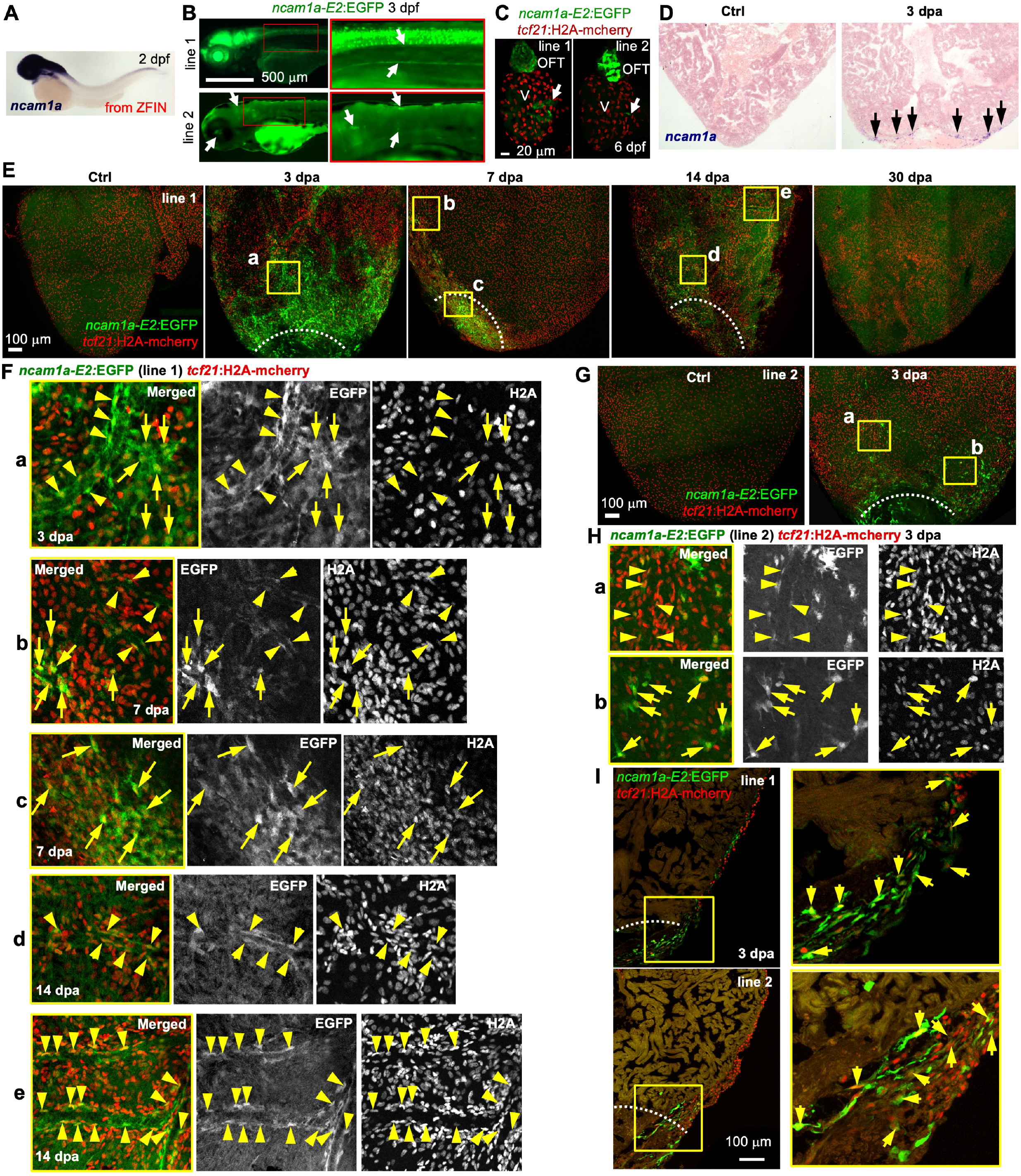
*ncam1a-E2* directs injury-induced epicardial gene expression. **(A)** Whole-mount *in situ* hybridization showing *ncam1a* expression in a 2 dpf embryo. Adapted from ZFIN (Thisse et al., 2008). **(B)** Larval expression of *ncam1a-E2:EGFP* lines at 3 dpf. Close-up views of the framed regions are shown on the right. Scale bar, 500 μm. **(C)** Whole-mount images of 6 dpf hearts showing EGFP expression in the outflow tract (OFT) and the atrioventricular valves (arrows). *tcf21*:H2A-mCherry (red) labels the epicardial cells. V, ventricle. Scale bar, 20 μm. **(D)** *In situ* hybridization results showing *ncam1a* expression on the ventricular surface around the injury site at 3 dpa (arrows). **(E)** Whole-mount images of the ventricular surface showing expressions of *ncam1a-E2:EGFP* line 1 (green) in uninjured (Ctrl) and 3, 7, 14, and 30 dpa samples. *tcf21*:H2A-mCherry (red) labels the epicardial cells. White dashed lines indicate the injury sites. The framed regions are enlarged to show details in (F). Scale bar, 100 μm. **(F)** Arrows or arrowheads indicate representative cells that express both EGFP (green) and *tcf21*:H2A-mCherry (red). Arrowheads in (a), (b), (d), and (e) denote *tcf21*^+^ perivascular cells. **(G)** Whole-mount images of the ventricular surface showing expressions of *ncam1a-E2:EGFP* line 2 (green) in uninjured (Ctrl) and 3 dpa samples. Magnified views of the framed region are shown in (H). **(H)** Single-channel images are shown in grayscale. Arrows and arrowheads (perivascular) indicate representative double-positive cells. **(I)** Section images demonstrating expressions of *ncam1a-E2:EGFP* (green) in *tcf21*:H2A-mCherry^+^ (red) cells. Magnified view of the framed regions is shown on the right. White dashed lines indicate the injury sites. Scale bar, 100 μm. Arrows denote representative double-positive cells.

We next characterized *ncam1a-E4*. Three *ncam1a-E4:EGFP* lines displayed very weak whole-body larval EGFP expression without clear specificity (Figure 7A, B). With an anti-EGFP antibody staining in adult hearts, we constantly observed a small population of EGFP^+^; *tcf21:*H2A-mCherry^+^ cells on the ventricular surface after injury (Figure 7C). Enhancers may reside upstream or downstream to the TSS of the regulated gene and may function together as a cluster to exert additive and synergistic actions (Choi et al., 2021; Hnisz et al., 2013). We asked whether putting multiple enhancers in series would enhance the activity. To this end, a second *E4* element was inserted after the poly-A signal of the *ncam1a-E4:EGFP* construct (Figure 7D), and one stable line was identified (*ncam1a-E4E4:EGFP*). *ncam1a-E4E4:EGFP* is expressed in the eye, tail, and heart muscles but not larval *tcf21*^+^ epicardial cells (Figure 7D, E). Adult heart EGFP expression was only observed upon heart injury. At 3 dpa, strong EGFP expression in *tcf21:*H2A-mCherry^+^ cells is located at the injury site, while slightly weaker expression was also observed in the entire epicardium distal to the injury site (Figure 7F, H), distinguishing its activity from the *ncam1a-E2:EGFP* lines. EGFP expression is reduced after 3 dpa, restricted to the injury site at 7 dpa, and undetectable at 30 dpa. Unlike *ncam1a-E2*, no perivascular cell expression was observed for either *ncam1a-E4E4:EGFP* or *ncam1a-E4:EGFP* lines. Following the same strategy, we asked whether combining two *ncam1a-E2* sequences would enhance its activity (Figure S5A). Compared to *ncam1a-E2:EGFP*, *ncam1a-E2E2:EGFP* lines have relatively similar EGFP expression in the embryos (Figure S5B, C). These *ncam1a-E2E2:EGFP* lines demonstrated apparently identical expression patterns and the same trend of expression levels as the *ncam1a-E2:EGFP* reporters (peak at 3 dpa, decrease through 30 dpa; Figure S5D-F). Although the larval expression was variable between lines of each enhancer, the adult epicardial expression was remarkably consistent across lines. These results suggest that n*cam1a-E2* and *-E4* are epicardial TREEs and likely regulate *ncam1a* expression during heart regeneration. Interestingly, their distinct expression dynamics indicate that individual enhancers may combine to generate an overall expression domain. This is comparable to the overlapping but distinct activities of multiple developmental enhancers linked to one gene (Dickel et al., 2018; Dunipace et al., 2019; Osterwalder et al., 2018), a regulatory strategy of which ostensibly applies to regeneration contexts.

**Figure 7.**
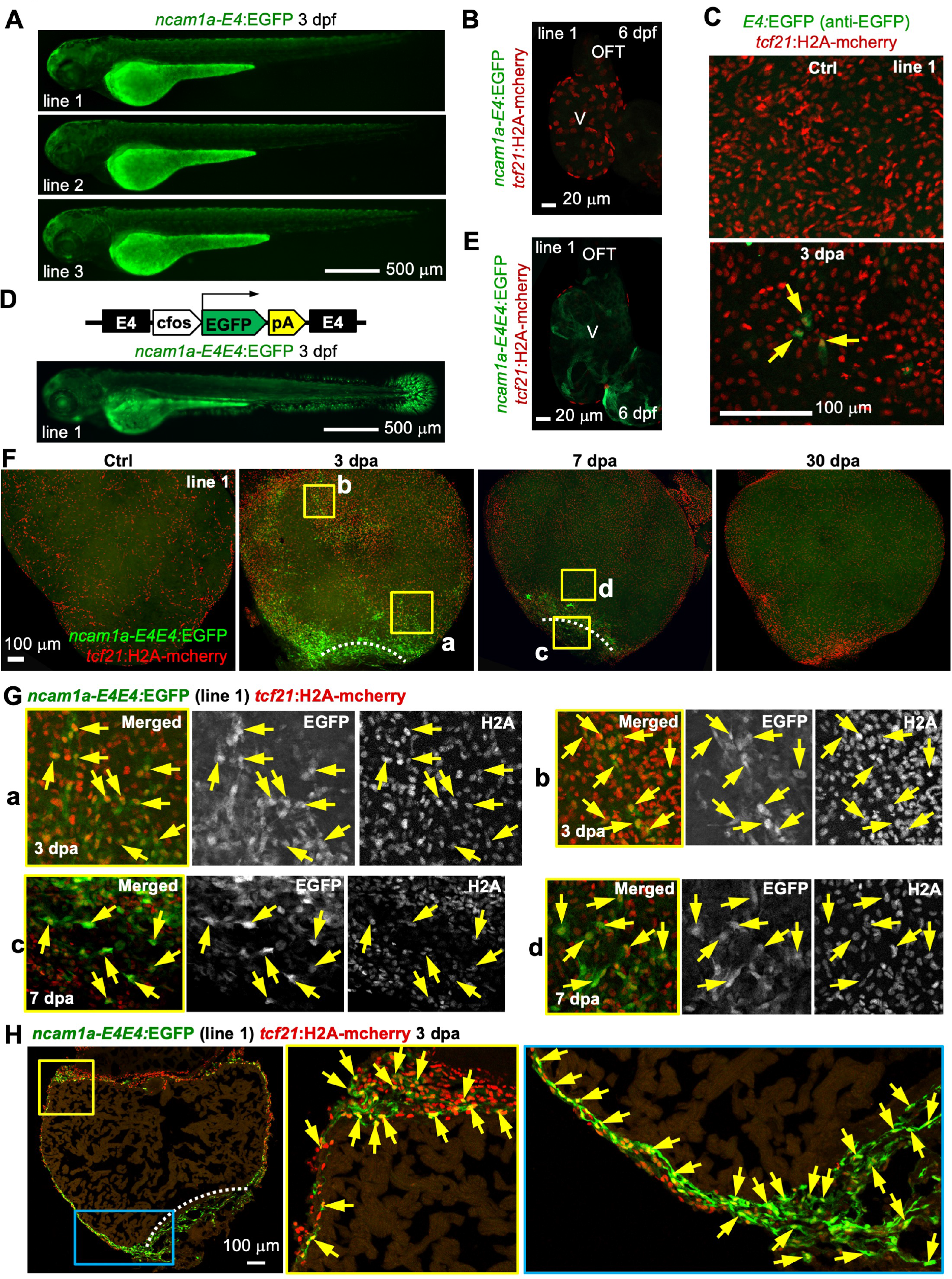
*ncam1a-E4* directs injury-induced epicardial gene expression. **(A)** Larval expression of *ncam1a-E4:EGFP* lines. Scale bar, 500 μm. **(B)** Whole-mount image of a 6 dpf hearts with *tcf21*:H2A-mCherry (red) labels the epicardial cells. No EGFP expression was observed. V, ventricle. Scale bar, 20 μm. **(C)** Whole-mount images of the ventricular surface showing expressions of an *ncam1a-E4:EGFP* line in uninjured (Ctrl) and 3 dpa sample. An anti-EGFP antibody staining was applied to detect the EGFP expression. **(D)** Top: the *ncam1a-E4E4:EGFP* reporter construct. Bottom: larval expression of *ncam1a-E4E4:EGFP* reporter line. Scale bar, 500 μm. **(E)** Optical section image of a 6 dpf heart (whole-mounted) showing GFP expression in the muscle. *tcf21*:H2A-mCherry (red) labels the epicardial cells. V, ventricle. Scale bar, 20 μm. **(F)** Whole-mount images of the ventricular surface showing expressions of *ncam1a-E4E4:EGFP* line 1 (green) in uninjured (Ctrl) and 3, 7, and 30 dpa samples. *tcf21*:H2A-mCherry (red) labels the epicardial cells. White dashed lines indicate the injury sites. The framed regions are enlarged to show details in (G). Scale bar, 100 μm. **(G)** Single-channel images are shown in grayscale. Arrows indicate representative GFP^+^mCherry^+^ cells. **(H)** Section images demonstrating expressions of *ncam1a-E4E4:EGFP* (green) in *tcf21*:H2A-mCherry^+^ (red) cells. Magnified view of the framed regions is shown on the right. White dashed lines indicate the injury sites. Arrows denote representative double-positive cells. Scale bar, 100 μm.

### Conclusions

The profiling we present can guide identification of factors that bind and are upstream of candidate and validated regulatory sequences, as well as target genes linked to these sequences, to help elucidate the epicardial injury response during heat regeneration. Huang et al. previously defined epicardial enhancers by analyzing the evolutionarily conserved regions linked to epicardial genes, including *Raldh2* and *Wt1,* and found enhancers that can direct expression both in developing hearts and in response to injury (Huang et al., 2012). By contrast, the epicardial TREEs we identified direct injury-induced but not developmental expression, suggesting they are customized to regeneration machinery. We found that multiple TREEs linked to a regeneration-regulated gene can possess either matching (*gnai3-E1, -E2*) or partially overlapping/complementary (*ncam1a-E2, -E4*) regulatory controls, suggesting additive, redundant, or synergetic effects of enhancers during regeneration. In this regard, deleting one of the enhancers for each gene may not fully abolish the injury-induced gene expression, which may explain why deletions of single TREEs typically can cause minor or no effect on gene expression (Thompson et al., 2020). We expect that the epicardial TREEs validated here, as well as putative TREEs implicated by the profiles, can be employed to generate transgenic lines that enable injury-induced expression of gene cassettes, for instance, to direct targeted expression of pro-regenerative factors (Bersell et al., 2009; Gemberling et al., 2015; Kang et al., 2016; Wei et al., 2015).

## MATERIALS AND METHODS

### Zebrafish maintenance and procedures

Adult zebrafish of the Ekkwill and Ekkwill/AB strains were maintained as described (Poss et al., 2002; Wang et al., 2011). Briefly, the water temperature was maintained at 28°C, and fish were kept on a 14/10 light/dark cycle at a density of 5-10 fish per liter. Animals of both sexes were used for adult experiments. Heart resection injury was done as described previously (Poss et al., 2002). Published transgenic lines used in this study were *Tg(tcf21:H2A-mCherry)^pd252^* (Cao et al., 2017) and *Tg(tcf21:nucEGFP)^pd41^* (Kikuchi et al., 2011). All transgenic strains were analyzed as hemizygotes. Animal procedures were approved by Animal Care and Use Committees at Duke University and Weill Cornell Medical College.

### Generation of transgenic reporters

Putative enhancer regions were amplified from genomic DNA using primers listed in Table S7 and inserted upstream of the 95 bp minimal mouse *c-fos* promoter directing EGFP (Fivaz et al., 2000). The entire *enhancer-fos-EGFP-SV40 poly A* cassette is flanked by two I-Sce meganuclease restriction sites that facilitate transgenesis (Babaryka et al., 2009). These constructs were injected into one-cell-stage wild-type embryos using standard meganuclease transgenesis techniques (Babaryka et al., 2009). To isolate stable lines, larvae were examined for EGFP expression or genotyped for *EGFP* insertions at 1-5 dpf. Twenty-one stable lines were established in this study (listed in Table S8).

### RNA sequencing and analysis

Partial ventricular resection was performed with *tcf21:nucEGFP* animals as described (Poss et al., 2002). Ventricles were collected at 3 and 7 dpa together with uninjured clutchmates. 30-50 ventricles per group from fish of mixed sexes were dissociated using Liberase DH (Roche) and EGFP^+^ epicardial cells were isolated by FACS as described previously (Cao et al., 2016). We collected 110-170k cells from each group of 2 biological replicates. Total RNA was purified using a Qiagen RNeasy Plus Micro Kit. The library was constructed using SMARTer Ultra Low Input RNA Kit for Sequencing, and sequencing was performed using an Illumina HiSeq 2000, with 82-94 million 50-bp single-end reads obtained for each library. Sequences were aligned to the zebrafish genome (danRer10) using TopHat2 (Kim et al., 2013). Transcript levels were quantified using HTseq (Anders et al., 2015). Differential expression analysis was done using the Bioconductor package DESeq2 (Love et al., 2014).

### ATAC sequencing and analysis

Partial ventricular resection was performed with *tcf21:nucEGFP* animals. Ventricles were collected at 3 and 7 dpa together with uninjured clutchmates. Four biological replicates were performed for each group. Ventricle dissociation and epicardial cells isolation were done as described above. ATAC-seq libraries were made from 50 - 198 k epicardial cells per sample as described previously (Buenrostro et al., 2013). Sequencing was performed using an Illumina HiSeq 2000 with 89 - 167 million 50-nt single reads obtained for each cell library.

ATAC-seq reads were trimmed for adaptors before aligning to the DanRer10 genome using Bowtie with two mismatches allowed and mapping up to 4 sites (Langmead et al., 2009). Reads mapped to mitochondrial DNA were removed, and alignments were filtered for PCR artifacts. Peaks were called using MACS v2.1.4 with p-value < 0.01 and shift -37 bp and extend 73 bp (Zhang et al., 2008). Normalized UCSC browser tracks were generated by conversion of bam format alignments to bp resolution bigWig files and scaling by the total number of mapped reads. The Bioconductor package DiffBind V3.0.15 was used for signal normalization and differential analysis (Ross-Innes et al., 2012). A fold change greater than two and a p-value of less than 0.05 were used to filter the significant differential peaks. Genomic distribution of peaks was analyzed using the Bioconductor package ChIPpeakAnno v3.27.2 (Zhu et al., 2010). Packages ChIPpeakAnno (Zhu et al., 2010) and complexHeatmap v2.6.2 (Gu et al., 2016) were used to annotate the peaks and generate heatmaps. The differential peaks were annotated by nearest gene start site to the center of peaks. ATAC-Seq peaks were paired to RNA-Seq differential expression data by annotated nearest gene symbols. Motif analysis was done by using HOMER v4.9.1 (Heinz et al., 2010). The conserved open-chromatin regions were identified by comparing our dataset with the published zebrafish conserved non-genic elements (zCNEs) database for at least 1 bp overlapped regions (Hiller et al., 2013). Ventricle Chip-seq (H3K27Ac) dataset was downloaded from Gene Expression Omnibus (GEO) under accession number GSE75894.

### Histology and microscopy

Images of fluorescence transgenes in live embryos or isolated hearts were captured using a Zeiss Axiozoom V16 microscope. Freshly collected hearts were fixed with 4% paraformaldehyde (PFA) for 1.5 h at room temperature or overnight at 4 degrees. Immunostaining of whole-mounted hearts was done as described previously (Cao et al., 2017; Wang et al., 2015). The primary antibody used in this study is mouse anti-EGFP (ThermoFisher, A11120). The secondary antibody used in this study was Alexa Fluor 488 goat anti-mouse (ThermoFisher, A11029). Fixed hearts were either mounted with Fluoromount G (Southern Biotechnology, 0100-01) between two coverslips (allowing imaging of both ventricular surfaces) or applied to cryosection at a 10 μm thickness. Whole-mounted and sectioned heart tissues were imaged used a Zeiss LSM800 confocal microscope.

### Data Availability

The RNA-seq and ATAC-seq datasets have been deposited at GEO under accession number GSE89444 (reviewers access token: cjaleeecfbenzgt).

## ACKNOWLEDGMENTS

We thank the Zebrafish Core Facility staff at Duke University and Weill Cornell for fish care, M. Harrison and J. Kang for comments on the manuscript. This work was supported by a Rudin Foundation Fellowship to Y.X., National Institutes of Health (NIH) grants (R01HL131319 and R35HL150713) and an American Heart Association (AHA) Merit Award to K.D.P., AHA Career Development Award (18CDA34110108), Weill Cornell Start-up fund, and NIH grant (R01HL155607) to J.C.

## AUTHORS CONTRIBUTIONS

Conceptualization, K.D.P. and J.C.; Methodology, G.C. and J.C.; Investigation, Y.C., Y.X., J.B.B., L.S., J.O., A.S., and T.C.; Resources, G.C., K.D.P., and J.C.; Writing and editing, Y.C., K.D.P., and J.C.; Funding Acquisition, Y.X., K.D.P., and J.C.

## Supplementary figures

**Figure S1.**
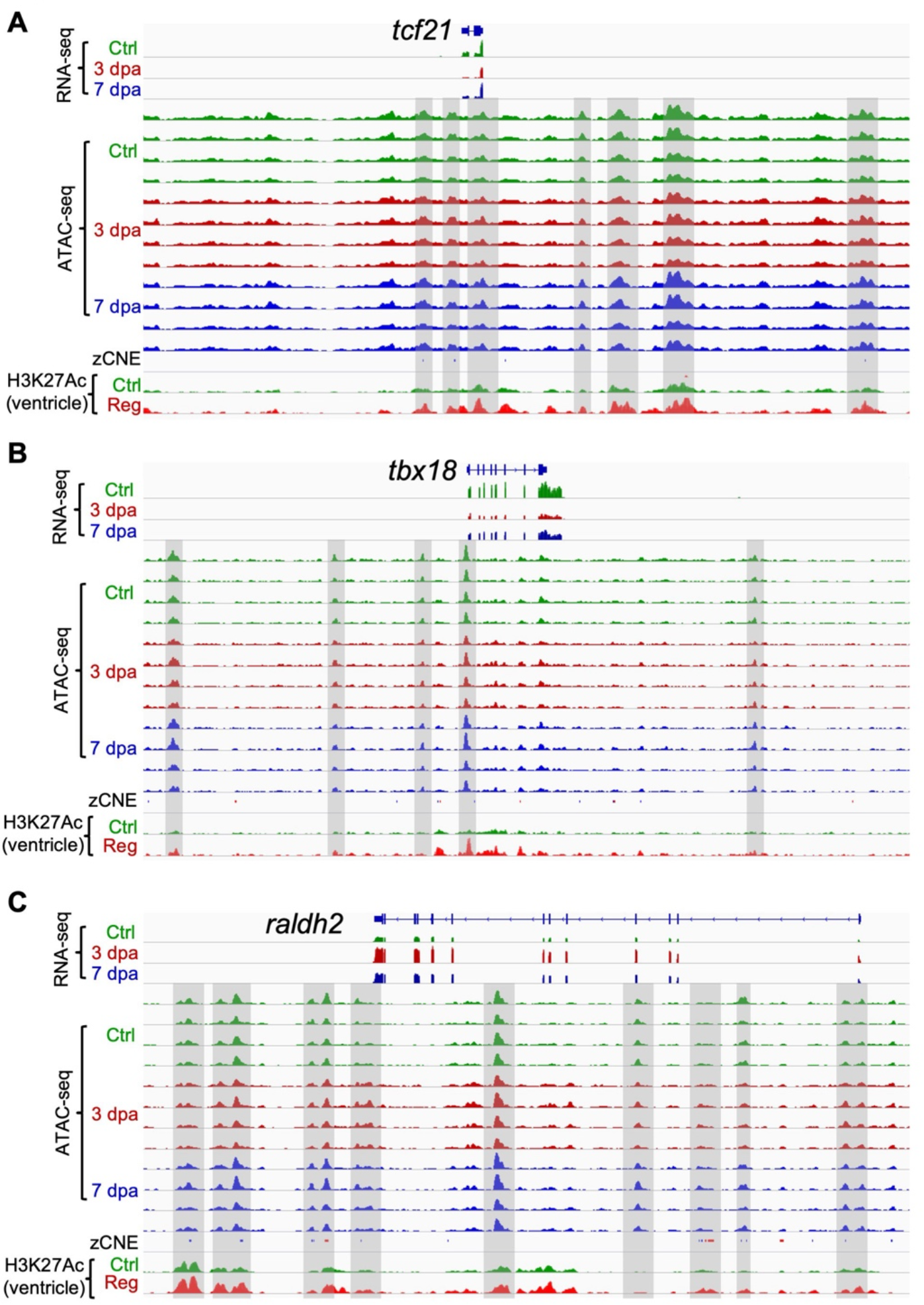
Browser tracks of the genomic region comprising *tcf21*, *tbx18*, or *raldh2*. Chromatin accessibility profiles in the epicardium, the zCNE sites, and the whole-ventricle H3K27Ac profile are shown. Selected putative enhancer regions are highlighted in gray boxes.

**Figure S2.**
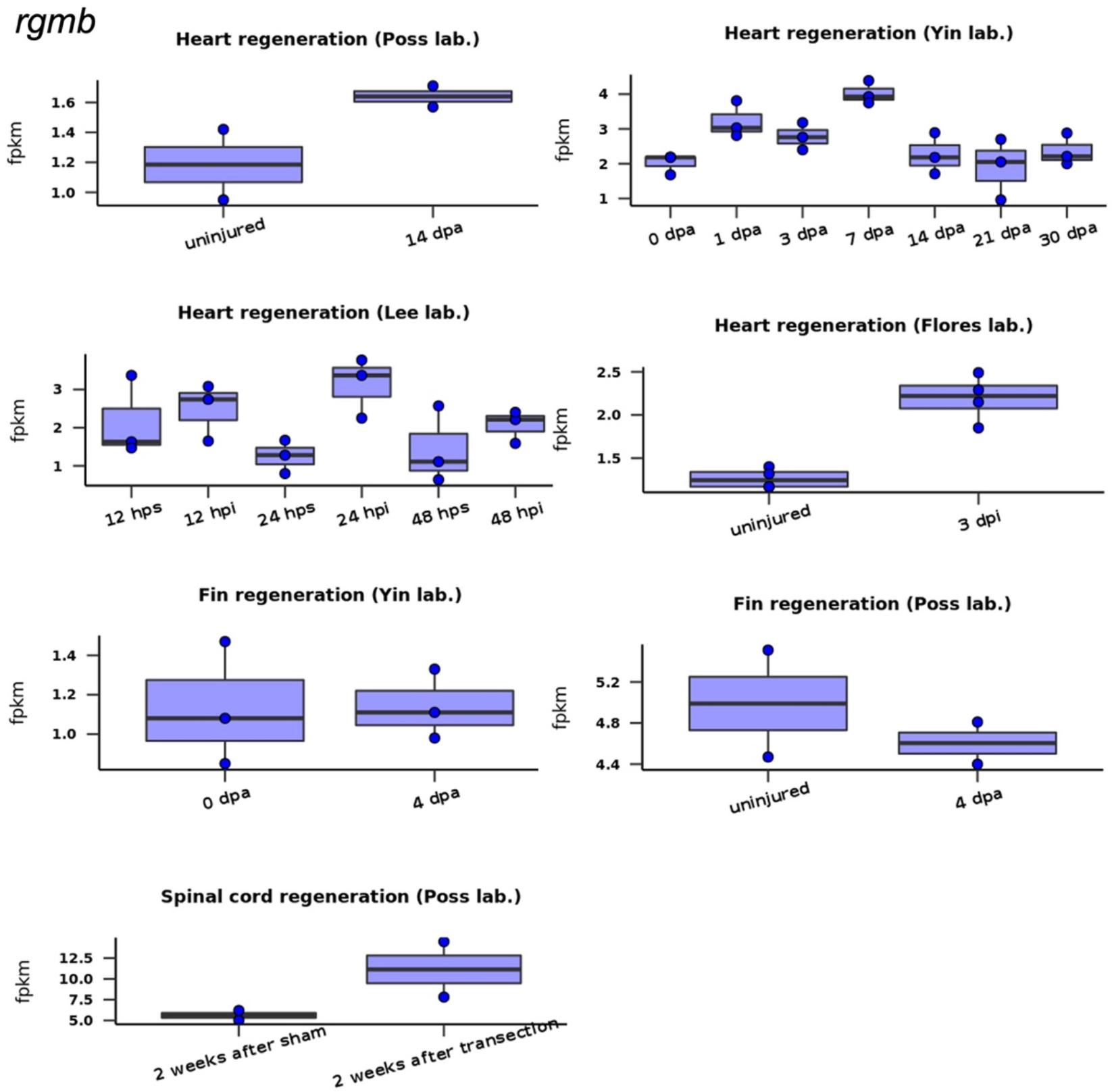
*rgmb* expression during zebrafish heart, fin, or spinal cord regeneration. Plots were generated using the Zebrafish Regeneration Database.

**Figure S3.**
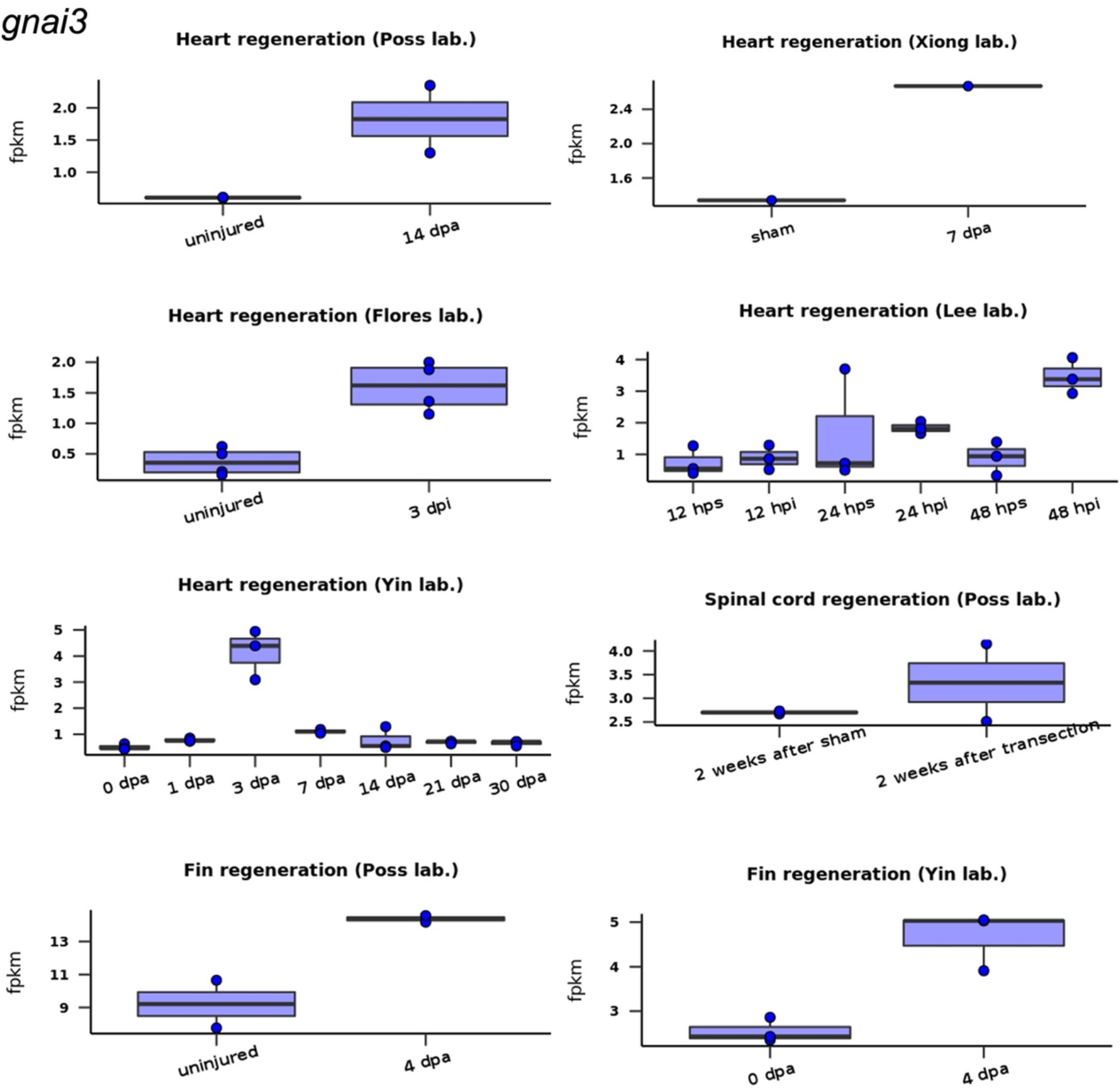
*gnai3* expression during zebrafish heart, fin, or spinal cord regeneration. Plots were generated using the Zebrafish Regeneration Database.

**Figure S4.**
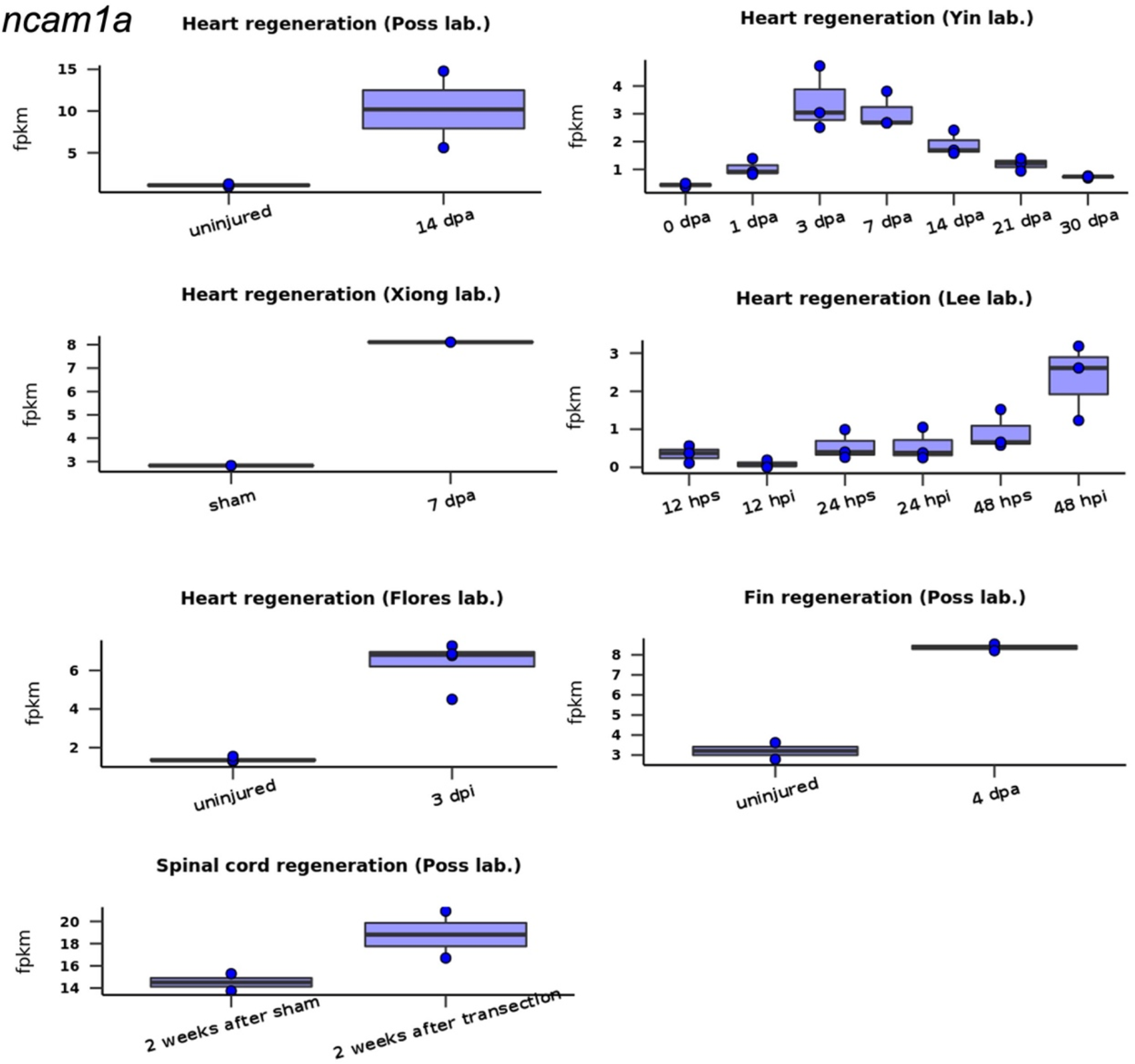
*ncam1a* expression during zebrafish heart, fin, or spinal cord regeneration. Plots were generated using the Zebrafish Regeneration Database.

**Figure S5.**
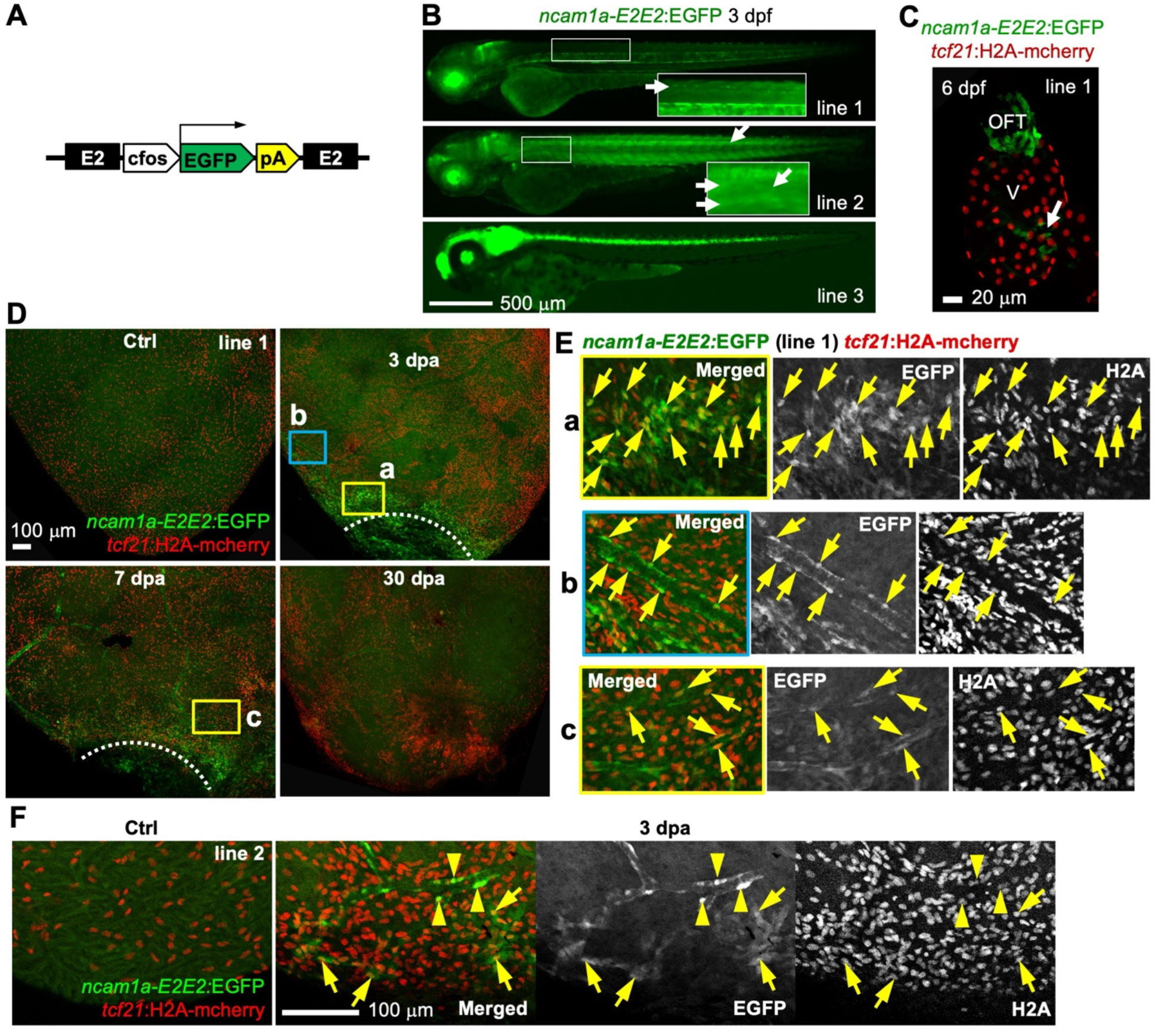
*ncam1a-E2E2* directs injury-induced epicardial gene expression. **(A)** The *ncam1a-E2E2:EGFP* enhancer reporter construct. A second *ncam1a-E2* enhancer is placed after the poly-A sequence. **(B)** Larval expression of the *ncam1a-E2E2:EGFP* line. Besides the eye, brain, spinal cord, and notochord expressions, additional EGFP signals are detected in the skeletal muscle for line 2. Scale bar, 500 μm. **(C)** Whole-mount images of a 6 dpf heart showing EGFP expression in the outflow tract (OFT) and the atrioventricular valves (arrows). *tcf21*:H2A-mCherry (red) labels the epicardial cells. V, ventricle. Scale bar, 20 μm. **(D)** Whole-mount images of the ventricular surface showing expressions of *ncam1a-E2E2:EGFP* line 1 (green) in uninjured (Ctrl) and 3, 7, and 30 dpa samples. *tcf21*:H2A-mCherry (red) labels the epicardial cells. White dashed lines indicate the injury sites. The framed regions are enlarged to show details in (E). Scale bar, 100 μm. **(E)** Single-channel images are shown in grayscale. Arrows indicate representative GFP^+^mCherry^+^ cells. Double-positive perivascular cells are shown in (b) and (c). **(F)** Whole-mount images showing expressions of *ncam1a-E2E2:EGFP* line 2 (green) in uninjured (Ctrl) and 3 dpa samples. *tcf21*:H2A-mCherry (red) labels the epicardial cells. White dashed lines indicate the injury sites. Single-channel images of the 3 dpa sample are shown in grayscale. Arrowheads (perivascular cells) and arrows indicate representative EGFP^+^mCherry^+^ cells. Scale bar, 100 μm.

## Supplementary Tables

**Table S1.** Alignment information of the ATAC-seq datasets.

**Table S2.** List of differential ATAC-seq peaks derived from paired comparisons.

**Table S3.** Enrichment annotation results of the ATAC-seq peaks with increased accessibility at 3 dpa.

**Table S4.** List of differential transcripts from paired comparisons.

**Table S5.** List of differential ATAC-seq peaks linked to nearby differential transcripts in 3 dpa versus uninjured (Ctrl) samples.

**Table S6.** List of upregulated ATAC-seq peaks (3 dpa) containing conserved sequences.

**Table S7.**
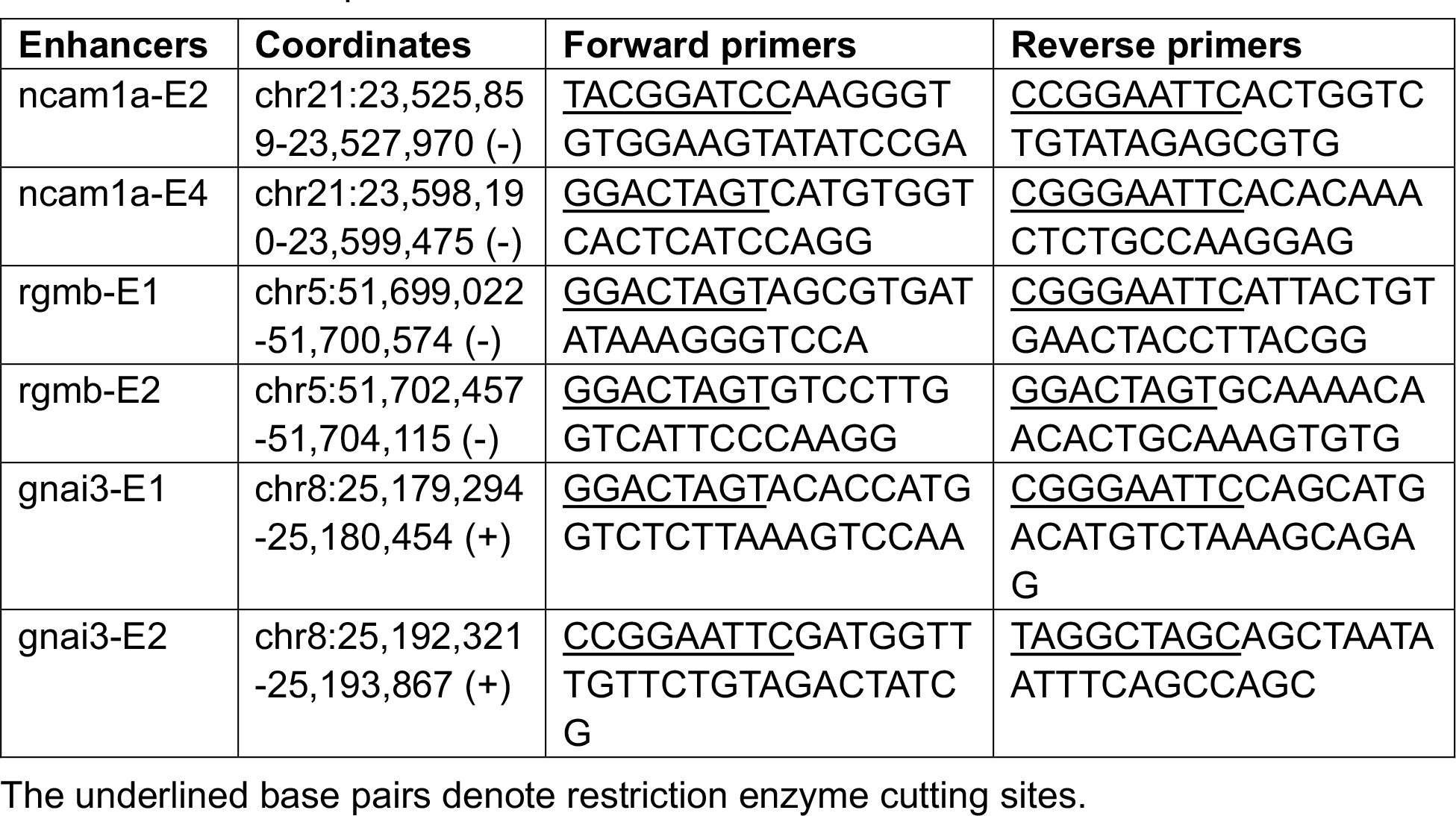
Primer sequences.

**Table S8.** Enhancer lines used in this study.

| # | Strain Name | Allele |
| --- | --- | --- |
| 1 | *Tg(ncam1a-E2:EGFP) line 1* | *wcm103* |
| 2 | *Tg(ncam1a-E2:EGFP) line 2* |  |
| 3 | *Tg(ncam1a-E2E2:EGFP) line 1* | *wcm104* |
| 4 | *Tg(ncam1a-E2E2:EGFP) line 2* |  |
| 5 | *Tg(ncam1a-E2E2:EGFP) line 3* |  |
| 6 | *Tg(ncam1a-E4:EGFP) line 1* | *wcm105* |
| 7 | *Tg(ncam1a-E4:EGFP) line 2* |  |
| 8 | *Tg(ncam1a-E4:EGFP) line 3* |  |
| 9 | *Tg(ncam1a-E4E4:EGFP) line 1* | *wcm106* |
| 10 | *Tg(rgmb-E1:EGFP) line 1* | *pd340* |
| 11 | *Tg(rgmb-E1:EGFP) line 2* |  |
| 12 | *Tg(rgmb-E2:EGFP) line 1* | *pd341* |
| 13 | *Tg(rgmb-E2:EGFP) line 2* |  |
| 14 | *Tg(rgmb-E2:EGFP) line 3* |  |
| 15 | *Tg(gnai3-E1:EGFP) line 1* | *pd342* |
| 16 | *Tg(gnai3-E1:EGFP) line 2* |  |
| 17 | *Tg(gnai3-E1:EGFP) line 3* |  |
| 18 | *Tg(gnai3-E2:EGFP) line 1* | *pd343* |
| 19 | *Tg(gnai3-E2:EGFP) line 2* |  |
| 20 | *Tg(gnai3-E2:EGFP) line 3* |  |
| 21 | *Tg(gnai3-E2:EGFP) line 4* |  |

## REFERENCES

1. Anders, S., Pyl, P. T. and Huber, W. (2015). HTSeq--a Python framework to work with high-throughput sequencing data. Bioinformatics 31, 166–169.

2. Babaryka, A., Kuhn, E. and Koster, R. W. (2009). In vivo synthesis of meganuclease for generating transgenic zebrafish Danio rerio. J Fish Biol 74, 452–457.

3. Begeman, I. J., Shin, K., Osorio-Mendez, D., Kurth, A., Lee, N., Chamberlain, T. J., Pelegri, F. J. and Kang, J. (2020). Decoding an organ regeneration switch by dissecting cardiac regeneration enhancers. Development 147.

4. Beisaw, A., Kuenne, C., Guenther, S., Dallmann, J., Wu, C. C., Bentsen, M., Looso, M. and Stainier, D. Y. R. (2020). AP-1 Contributes to Chromatin Accessibility to Promote Sarcomere Disassembly and Cardiomyocyte Protrusion During Zebrafish Heart Regeneration. Circ Res 126, 1760–1778.

5. Bersell, K., Arab, S., Haring, B. and Kuhn, B. (2009). Neuregulin1/ErbB4 signaling induces cardiomyocyte proliferation and repair of heart injury. Cell 138, 257–270.

6. Buenrostro, J. D., Giresi, P. G., Zaba, L. C., Chang, H. Y. and Greenleaf, W. J. (2013). Transposition of native chromatin for fast and sensitive epigenomic profiling of open chromatin, DNA-binding proteins and nucleosome position. Nature methods 10, 1213–1218.

7. Cao, J., Navis, A., Cox, B. D., Dickson, A. L., Gemberling, M., Karra, R., Bagnat, M. and Poss, K. D. (2016). Single epicardial cell transcriptome sequencing identifies Caveolin 1 as an essential factor in zebrafish heart regeneration. Development 143, 232–243.

8. Cao, J. and Poss, K. D. (2018). The epicardium as a hub for heart regeneration. Nat Rev Cardiol 15, 631– 647.

9. Cao, J., Wang, J., Jackman, C. P., Cox, A. H., Trembley, M. A., Balowski, J. J., Cox, B. D., De Simone, A., Dickson, A. L., Di Talia, S., et al. (2017). Tension Creates an Endoreplication Wavefront that Leads Regeneration of Epicardial Tissue. Dev Cell 42, 600–615 e604.

10. Chablais, F. and Jazwinska, A. (2012). The regenerative capacity of the zebrafish heart is dependent on TGFbeta signaling. Development 139, 1921–1930.

11. Chen, A., Han, Y. and Poss, K. D. (2020). Regulation of zebrafish fin regeneration by vitamin D signaling. Developmental dynamics : an official publication of the American Association of Anatomists.

12. Choi, J., Lysakovskaia, K., Stik, G., Demel, C., Soding, J., Tian, T. V., Graf, T. and Cramer, P. (2021). Evidence for additive and synergistic action of mammalian enhancers during cell fate determination. eLife 10.

13. Choi, W. Y., Gemberling, M., Wang, J., Holdway, J. E., Shen, M. C., Karlstrom, R. O. and Poss, K. D. (2013). In vivo monitoring of cardiomyocyte proliferation to identify chemical modifiers of heart regeneration. Development 140, 660–666.

14. Dickel, D. E., Ypsilanti, A. R., Pla, R., Zhu, Y., Barozzi, I., Mannion, B. J., Khin, Y. S., Fukuda-Yuzawa, Y., Plajzer-Frick, I., Pickle, C. S., et al. (2018). Ultraconserved Enhancers Are Required for Normal Development. Cell 172, 491–499 e415.

15. Duncan, B. W., Murphy, K. E. and Maness, P. F. (2021). Molecular Mechanisms of L1 and NCAM Adhesion Molecules in Synaptic Pruning, Plasticity, and Stabilization. Front Cell Dev Biol 9, 625340.

16. Dunipace, L., Akos, Z. and Stathopoulos, A. (2019). Coacting enhancers can have complementary functions within gene regulatory networks and promote canalization. PLoS genetics 15, e1008525.

17. Feng, T., Meng, J., Kou, S., Jiang, Z., Huang, X., Lu, Z., Zhao, H., Lau, L. F., Zhou, B. and Zhang, H. (2019). CCN1-Induced Cellular Senescence Promotes Heart Regeneration. Circulation 139, 2495–2498.

18. Fivaz, J., Bassi, M. C., Pinaud, S. and Mirkovitch, J. (2000). RNA polymerase II promoter-proximal pausing upregulates c-fos gene expression. Gene 255, 185–194.

19. Frankel, N., Davis, G. K., Vargas, D., Wang, S., Payre, F. and Stern, D. L. (2010). Phenotypic robustness conferred by apparently redundant transcriptional enhancers. Nature 466, 490–493.

20. Gemberling, M., Karra, R., Dickson, A. L. and Poss, K. D. (2015). Nrg1 is an injury-induced cardiomyocyte mitogen for the endogenous heart regeneration program in zebrafish. eLife 4.

21. Goldman, J. A., Kuzu, G., Lee, N., Karasik, J., Gemberling, M., Foglia, M. J., Karra, R., Dickson, A. L., Sun, F., Tolstorukov, M. Y., et al. (2017). Resolving Heart Regeneration by Replacement Histone Profiling. Dev Cell 40, 392–404 e395.

22. Gonzalez-Rosa, J. M., Peralta, M. and Mercader, N. (2012). Pan-epicardial lineage tracing reveals that epicardium derived cells give rise to myofibroblasts and perivascular cells during zebrafish heart regeneration. Developmental biology 370, 173–186.

23. Gu, Z., Eils, R. and Schlesner, M. (2016). Complex heatmaps reveal patterns and correlations in multidimensional genomic data. Bioinformatics 32, 2847–2849.

24. Harris, R. E., Setiawan, L., Saul, J. and Hariharan, I. K. (2016). Localized epigenetic silencing of a damage-activated WNT enhancer limits regeneration in mature Drosophila imaginal discs. eLife 5.

25. Heinz, S., Benner, C., Spann, N., Bertolino, E., Lin, Y. C., Laslo, P., Cheng, J. X., Murre, C., Singh, H. and Glass, C. K. (2010). Simple combinations of lineage-determining transcription factors prime cis-regulatory elements required for macrophage and B cell identities. Mol Cell 38, 576–589.

26. Hiller, M., Agarwal, S., Notwell, J. H., Parikh, R., Guturu, H., Wenger, A. M. and Bejerano, G. (2013). Computational methods to detect conserved non-genic elements in phylogenetically isolated genomes: application to zebrafish. Nucleic acids research 41, e151.

27. Hnisz, D., Abraham, B. J., Lee, T. I., Lau, A., Saint-Andre, V., Sigova, A. A., Hoke, H. A. and Young, R. A. (2013). Super-enhancers in the control of cell identity and disease. Cell 155, 934–947.

28. Hornblad, A., Bastide, S., Langenfeld, K., Langa, F. and Spitz, F. (2021). Dissection of the Fgf8 regulatory landscape by in vivo CRISPR-editing reveals extensive intra- and inter-enhancer redundancy. Nat Commun 12, 439.

29. Hu, H., Lin, S., Wang, S. and Chen, X. (2020). The Role of Transcription Factor 21 in Epicardial Cell Differentiation and the Development of Coronary Heart Disease. Front Cell Dev Biol 8, 457.

30. Huang, G. N., Thatcher, J. E., McAnally, J., Kong, Y., Qi, X., Tan, W., DiMaio, J. M., Amatruda, J. F., Gerard, R. D., Hill, J. A., et al. (2012). C/EBP transcription factors mediate epicardial activation during heart development and injury. Science 338, 1599–1603.

31. Hui, S. P., Sheng, D. Z., Sugimoto, K., Gonzalez-Rajal, A., Nakagawa, S., Hesselson, D. and Kikuchi, K. (2017). Zebrafish Regulatory T Cells Mediate Organ-Specific Regenerative Programs. Dev Cell 43, 659–672 e655.

32. Kang, J., Hu, J., Karra, R., Dickson, A. L., Tornini, V. A., Nachtrab, G., Gemberling, M., Goldman, J. A., Black, B. L. and Poss, K. D. (2016). Modulation of tissue repair by regeneration enhancer elements. Nature 532, 201–206.

33. Karra, R., Foglia, M. J., Choi, W. Y., Belliveau, C., DeBenedittis, P. and Poss, K. D. (2018). Vegfaa instructs cardiac muscle hyperplasia in adult zebrafish. Proc Natl Acad Sci U S A 115, 8805–8810.

34. Kikuchi, K., Gupta, V., Wang, J., Holdway, J. E., Wills, A. A., Fang, Y. and Poss, K. D. (2011). tcf21+ epicardial cells adopt non-myocardial fates during zebrafish heart development and regeneration. Development 138, 2895–2902.

35. Kim, D., Pertea, G., Trapnell, C., Pimentel, H., Kelley, R. and Salzberg, S. L. (2013). TopHat2: accurate alignment of transcriptomes in the presence of insertions, deletions and gene fusions. Genome Biol 14, R36.

36. Kvon, E. Z., Waymack, R., Gad, M. and Wunderlich, Z. (2021). Enhancer redundancy in development and disease. Nat Rev Genet 22, 324–336.

37. Langmead, B., Trapnell, C., Pop, M. and Salzberg, S. L. (2009). Ultrafast and memory-efficient alignment of short DNA sequences to the human genome. Genome Biol 10, R25.

38. Lee, H. J., Hou, Y., Chen, Y., Dailey, Z. Z., Riddihough, A., Jang, H. S., Wang, T. and Johnson, S. L. (2020). Regenerating zebrafish fin epigenome is characterized by stable lineage-specific DNA methylation and dynamic chromatin accessibility. Genome Biol 21, 52.

39. Lepilina, A., Coon, A. N., Kikuchi, K., Holdway, J. E., Roberts, R. W., Burns, C. G. and Poss, K. D. (2006). A dynamic epicardial injury response supports progenitor cell activity during zebrafish heart regeneration. Cell 127, 607–619.

40. Liu, J., Wang, W., Liu, M., Su, L., Zhou, H., Xia, Y., Ran, J., Lin, H. Y. and Yang, B. (2016). Repulsive guidance molecule b inhibits renal cyst development through the bone morphogenetic protein signaling pathway. Cell Signal 28, 1842–1851.

41. Love, M. I., Huber, W. and Anders, S. (2014). Moderated estimation of fold change and dispersion for RNA-seq data with DESeq2. Genome Biol 15, 550.

42. Lu, H. and Huang, H. (2011). FOXO1: a potential target for human diseases. Curr Drug Targets 12, 1235– 1244.

43. Marin-Juez, R., El-Sammak, H., Helker, C. S. M., Kamezaki, A., Mullapuli, S. T., Bibli, S. I., Foglia, M. J., Fleming, I., Poss, K. D. and Stainier, D. Y. R. (2019). Coronary Revascularization During Heart Regeneration Is Regulated by Epicardial and Endocardial Cues and Forms a Scaffold for Cardiomyocyte Repopulation. Dev Cell 51, 503–515 e504.

44. Masters, M. and Riley, P. R. (2014). The epicardium signals the way towards heart regeneration. Stem cell research 13, 683–692.

45. Nakae, J., Kitamura, T., Silver, D. L. and Accili, D. (2001). The forkhead transcription factor Foxo1 (Fkhr) confers insulin sensitivity onto glucose-6-phosphatase expression. The Journal of clinical investigation 108, 1359–1367.

46. Nieto-Arellano, R. and Sanchez-Iranzo, H. (2019). zfRegeneration: a database for gene expression profiling during regeneration. Bioinformatics 35, 703–705.

47. Osterwalder, M., Barozzi, I., Tissieres, V., Fukuda-Yuzawa, Y., Mannion, B. J., Afzal, S. Y., Lee, E. A., Zhu, Y., Plajzer-Frick, I., Pickle, C. S., et al. (2018). Enhancer redundancy provides phenotypic robustness in mammalian development. Nature 554, 239–243.

48. Pennacchio, L. A., Bickmore, W., Dean, A., Nobrega, M. A. and Bejerano, G. (2013). Enhancers: five essential questions. Nat Rev Genet 14, 288–295.

49. Pfefferli, C. and Jazwinska, A. (2017). The careg element reveals a common regulation of regeneration in the zebrafish myocardium and fin. Nat Commun 8, 15151.

50. Poss, K. D., Wilson, L. G. and Keating, M. T. (2002). Heart regeneration in zebrafish. Science 298, 2188– 2190.

51. Ross-Innes, C. S., Stark, R., Teschendorff, A. E., Holmes, K. A., Ali, H. R., Dunning, M. J., Brown, G. D., Gojis, O., Ellis, I. O., Green, A. R., et al. (2012). Differential oestrogen receptor binding is associated with clinical outcome in breast cancer. Nature 481, 389–393.

52. Samad, T. A., Rebbapragada, A., Bell, E., Zhang, Y., Sidis, Y., Jeong, S. J., Campagna, J. A., Perusini, S., Fabrizio, D. A., Schneyer, A. L., et al. (2005). DRAGON, a bone morphogenetic protein co-receptor. The Journal of biological chemistry 280, 14122–14129.

53. Sarig, R., Rimmer, R., Bassat, E., Zhang, L., Umansky, K. B., Lendengolts, D., Perlmoter, G., Yaniv, K. and Tzahor, E. (2019). Transient p53-Mediated Regenerative Senescence in the Injured Heart. Circulation 139, 2491–2494.

54. Siles, A. M., Martinez-Hernandez, E., Araque, J., Diaz-Manera, J., Rojas-Garcia, R., Gallardo, E., Illa, I., Graus, F. and Querol, L. (2018). Antibodies against cell adhesion molecules and neural structures in paraneoplastic neuropathies. Ann Clin Transl Neurol 5, 559–569.

55. Simoes, F. C. and Riley, P. R. (2018). The ontogeny, activation and function of the epicardium during heart development and regeneration. Development 145.

56. Sugimoto, K., Hui, S. P., Sheng, D. Z. and Kikuchi, K. (2017). Dissection of zebrafish shha function using site-specific targeting with a Cre-dependent genetic switch. eLife 6.

57. Syrovatkina, V., Alegre, K. O., Dey, R. and Huang, X. Y. (2016). Regulation, Signaling, and Physiological Functions of G-Proteins. J Mol Biol 428, 3850–3868.

58. Thisse, B., Wright, G. J. and Thisse, C. (2008). Embryonic and Larval Expression Patterns from a Large Scale Screening for Novel Low Affinity Extracellular Protein Interactions. ZFIN Direct Data Submission (http://zfin.org).

59. Thompson, J. D., Ou, J., Lee, N., Shin, K., Cigliola, V., Song, L., Crawford, G. E., Kang, J. and Poss, K. D. (2020). Identification and requirements of enhancers that direct gene expression during zebrafish fin regeneration. Development 147.

60. Umer, H. M., Smolinska-Garbulowska, K., Marzouka, N.-a.-d., Khaliq, Z., Wadelius, C. and Komorowski, J. (2019). funMotifs: Tissue-specific transcription factor motifs. bioRxiv, 683722.

61. van Duijvenboden, K., de Bakker, D. E. M., Man, J. C. K., Janssen, R., Gunthel, M., Hill, M. C., Hooijkaas, I. B., van der Made, I., van der Kraak, P. H., Vink, A., et al. (2019). Conserved NPPB+ Border Zone Switches From MEF2- to AP-1-Driven Gene Program. Circulation 140, 864–879.

62. Wang, J., Cao, J., Dickson, A. L. and Poss, K. D. (2015). Epicardial regeneration is guided by cardiac outflow tract and Hedgehog signalling. Nature 522, 226–230.

63. Wang, J., Karra, R., Dickson, A. L. and Poss, K. D. (2013). Fibronectin is deposited by injury-activated epicardial cells and is necessary for zebrafish heart regeneration. Developmental biology 382, 427– 435.

64. Wang, J., Panakova, D., Kikuchi, K., Holdway, J. E., Gemberling, M., Burris, J. S., Singh, S. P., Dickson, A. L., Lin, Y. F., Sabeh, M. K., et al. (2011). The regenerative capacity of zebrafish reverses cardiac failure caused by genetic cardiomyocyte depletion. Development 138, 3421–3430.

65. Wang, W., Hu, C. K., Zeng, A., Alegre, D., Hu, D., Gotting, K., Ortega Granillo, A., Wang, Y., Robb, S., Schnittker, R., et al. (2020). Changes in regeneration-responsive enhancers shape regenerative capacities in vertebrates. Science 369.

66. Wang, Y., Zhou, Y. and Graves, D. T. (2014). FOXO transcription factors: their clinical significance and regulation. Biomed Res Int 2014, 925350.

67. Wei, K., Serpooshan, V., Hurtado, C., Diez-Cunado, M., Zhao, M., Maruyama, S., Zhu, W., Fajardo, G., Noseda, M., Nakamura, K., et al. (2015). Epicardial FSTL1 reconstitution regenerates the adult mammalian heart. Nature 525, 479–485.

68. Wu, C. C., Kruse, F., Vasudevarao, M. D., Junker, J. P., Zebrowski, D. C., Fischer, K., Noel, E. S., Grun, D., Berezikov, E., Engel, F. B., et al. (2016). Spatially Resolved Genome-wide Transcriptional Profiling Identifies BMP Signaling as Essential Regulator of Zebrafish Cardiomyocyte Regeneration. Dev Cell 36, 36–49.

69. Xiao, Y., Hill, M. C., Zhang, M., Martin, T. J., Morikawa, Y., Wang, S., Moise, A. R., Wythe, J. D. and Martin, J. F. (2018). Hippo Signaling Plays an Essential Role in Cell State Transitions during Cardiac Fibroblast Development. Dev Cell 45, 153–169 e156.

70. Xiong, N., Kang, C. and Raulet, D. H. (2002). Redundant and unique roles of two enhancer elements in the TCRgamma locus in gene regulation and gammadelta T cell development. Immunity 16, 453– 463.

71. Yan, C., Xu, Z. and Huang, W. (2021). Cellular Senescence Affects Cardiac Regeneration and Repair in Ischemic Heart Disease. Aging Dis 12, 552–569.

72. Zhang, Y., Liu, T., Meyer, C. A., Eeckhoute, J., Johnson, D. S., Bernstein, B. E., Nusbaum, C., Myers, R. M., Brown, M., Li, W., et al. (2008). Model-based analysis of ChIP-Seq (MACS). Genome Biol 9, R137.

73. Zhu, L. J., Gazin, C., Lawson, N. D., Pages, H., Lin, S. M., Lapointe, D. S. and Green, M. R. (2010). ChIPpeakAnno: a Bioconductor package to annotate ChIP-seq and ChIP-chip data. BMC bioinformatics 11, 237.

